# PhaseTypeR: phase-type distributions in R with reward transformations and a view towards population genetics

**DOI:** 10.1101/2022.06.16.496381

**Authors:** Iker Rivas-González, Lars Nørvang Andersen, Asger Hobolth

**Affiliations:** Bioinformatics Research Center, Aarhus University, 8000 Aarhus C, Denmark

**Author notes:** . URL: https://github.com/rivasiker/.

**Keywords:** Ancestral process, coalescent theory, phase-type distributions, population genetics, **PhaseTypeR**

## Abstract

Phase-type distributions are a general class of models that are traditionally used in actuarial sciences and queuing theory, and more recently in population genetics. A phase-type distributed random variable is the time to absorption in a discrete or continuous time Markov chain on a finite state space with an absorbing state. The R package **PhaseTypeR** contains all the key functions—mean, (co)variance, probability density function, cumulative distribution function, quantile function, random sampling and reward transformations—for both continuous (PH) and discrete (DPH) phase-type distributions. Additionally, we have also implemented the multivariate continuous case (MPH) and the multivariate discrete case (MDPH). We illustrate the usage of **PhaseTypeR** in simple examples from population genetics (e.g. the time until the most recent common ancestor or the total number of mutations in an alignment of homologous DNA sequences), and we demonstrate the power of **PhaseTypeR** in more involved applications from population genetics, such as the coalescent with recombination and the structured coalescent. The multivariate distributions and ability to reward-transform are particularly important in population genetics, and a unique feature of **PhaseTypeR**.

## 1. Introduction

Phase-type distributions describe the time until absorption of a continuous or discrete-time Markov chain (Bladt and Nielsen 2017). The probabilistic properties of phase-type distributions (i.e. the probability density function, cumulative distribution function, quantile function, moments and generating functions) are well-described and analytically tractable using matrix manipulations.

Here we present **PhaseTypeR**, an R (R Core Team 2021) package that provides general-use core functions for continuous and discrete phase-type distributions, both for the univariate and the multivariate cases. **PhaseTypeR** can also be used to simulate from the underlying Markov chain of the phase-type objects. Additionally, the package allows for the reward transformation of phase-type distributions. The functions and objects in the package are intuitive and of general use, which enables the users to easily adapt them to their needs. **PhaseTypeR** is available on CRAN (https://CRAN.R-project.org/package=PhaseTypeR), and its documentation can be accessed through https://rivasiker.github.io/PhaseTypeR/.

The R packages already available for phase-type distributions are mainly tailored to applications in actuarial sciences and risk theory. In these cases, failure times or lifetimes are measured, and the corresponding phase-type distribution is estimated. We briefly summarize the software tools for phase-type distributions, their status and their main purpose:

- Christophe Dutang, Vincent Goulet and Mathieu Pigeon have contributed the R package **actuar** for the actuarial sciences (Dutang, Goulet, and Pigeon 2008), and the package is still under active development. The univariate continuous phase-type distribution is covered in terms of the density, cumulative distribution, moments and moment generating function (see https://CRAN.R-project.org/package=actuar).
- Louis Aslett has released an R package called **PhaseType** (Aslett and Wilson 2011; Aslett 2012), which is tailored to the problem of estimating a continuous phase-type distribution from failure times, and is an extension of a Markov Chain Monte Carlo algorithm developed by Bladt, Gonzalez, and Lauritzen (2003). However, the package is not maintained anymore, and it has been removed from CRAN (https://CRAN.R-project.org/package=PhaseType).
- Hiroyuki Okamura’s **mapfit** R package is concerned with fitting phase-type distributions of failure times in reliability systems (https://CRAN.R-project.org/package=mapfit, Okamura 2015; Okamura and Dohi 2015, 2016). Here, the parameters in a phase-type distribution are fitted using maximum likelihood estimation. The package can also fit a phase-type distribution from a probability density function.
- Martin Bladt and Jorge Yslas’s recent R package **matrixdist** (https://CRAN.R-project.org/package=matrixdist, Bladt and Yslas 2021) fits inhomogeneous phase-type (IPH) distributions (Albrecher and Bladt 2019). The EM-algorithm is used to estimate the parameters in the model. In Albrecher, Bladt, and Yslas (2020, advance online publication), an IPH distribution is fitted to the lifetimes of the Danish population that died in the year 2000 at ages 50 to 100. In the special homogeneous case of the IPH distributions, the package also provides the density, cumulative distribution function, quantile function, moments and opportunity of simulating from the distribution. Our package has the same features for the PH distribution, and additionally we consider reward transformations and the multivariate phase-type (MPH) extension. Furthermore, we provide the same functionality for the class of discrete phase-type (DPH) and multivariate discrete phase-type (MDPH) distributions.

Our implementation of phase-type functions in **PhaseTypeR** is of general use and not restricted to actuarial sciences and risk theory. Moreover, unlike the packages described above, **PhaseTypeR** includes reward transformations, the multivariate extensions of phase-type distributions, and both the discrete and continuous versions.

In this paper we exemplify the applications of the **PhaseTypeR** functions using several quantities in population genetics and, in particular, coalescent theory. More specifically, the time to the most recent common ancestor *T*_MRCA_, the total tree length *T*_total_ and the total branch lengths that give rise to e.g. singletons or doubletons are examples of continuous phase-type distributed variables, and viewed together they are continuous multivariate phase-type distributed (Hobolth, Siri-Jegousse, and Bladt 2019). Additionally, the individual elements of the site frequency spectrum are examples of discrete phase-type distributed variables, and the full site frequency spectrum is multivariate discrete phase-type distributed (Hobolth, Bladt, and Andersen 2021). These statements hold for the standard coalescent process, but also for more general time-homogeneous coalescent models such as the structured coalescent (Wakeley 2009, Section 5), the multiple merger coalescent (Tellier and Lemaire 2014), and the coalescent with recombination (Wakeley 2009, Section 7.2).

This paper presents the basic theory for phase-type distributions and demonstrates how to apply the distributions using **PhaseTypeR**. The paper is organized as follows. Section 2 is concerned with the univariate and multivariate continuous phase-type distributions. The basic phase-type object contains a subintensity matrix and an initial probability vector (potentially with a defect), and the four commonly associated functions `dPH` (probability density function), `pPH` (cumulative distribution function), `qPH` (quantile function) and `rPH` (random sampling). The function `rFullPH` provides a simulation of the full sample path from a continuous phasetype distribution. We then introduce the reward transformations and the multivariate continuous phase-type distribution. Section 3 follows the same structure and introduces the same type of functions (called `dDPH`, `pDPH`, `qDPH`, `rDPH`, `rFullDPH`) for the univariate and multivariate discrete phase-type distributions. In Section 4 we apply phase-type theory to understand the ancestral recombination graph for two loci and two samples, and in Section 5 we demonstrate how phase-type theory can be used to learn about a structured population. The paper ends with a conclusion and a discussion of future extensions and applications of our package.

## 2. Continuous phase-type distributions

### 2.1. Theory for the phase-type distribution

A continuous phase-type distribution is a sum of exponential distributions that occur sequentially until absorption. More specifically, a phase-type distribution is the time to absorption of a Markov jump process.

Following the notation in Bladt and Nielsen (2017), let {*X*_*t*_}_*t*≥0_ be a Markov jump process with *p* transient states and a single absorbing state. The time until absorption *τ* of such a process then follows a continuous phase-type distribution, where the rate matrix of the underlying Markov jump process **Λ** is given as

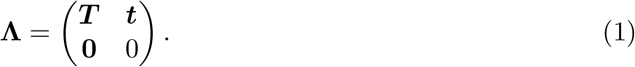

The *p* × *p* matrix ***T*** is the subintensity matrix, and the elements are the transition rates between the transient states. Because of the properties of rate matrices, all the rows of **Λ** sum to 0 (i.e. **Λ*e*** = **0**, where ***e*** is a vector of ones), which means that the phase-type distribution can be defined by the subintensity matrix ***T***, while the exit rate column vector ***t*** is given by ***t*** = −***T e***. Additionally, we have a vector of size *p* corresponding to the initial probabilities ***π***, such that *τ* ∼ PH(***π, T***) (i.e. *P* (*X*_0_ = *i*) = *π*_*i*_, *i* = 1, …, *p*). The sum of the initial probabilities (***πe***) might sum to less than 1. If this is the case, then we can define the defect as 1 − ***πe***, which corresponds to the probability of starting in the absorbing state without passing through any of the transient states (i.e. *P* (*X*_0_ = *p* + 1) = 1 − ***πe***).

The properties of phase-type distributions can easily be calculated due to their matrix-form representation. The mean, variance, probability density function and cumulative distribution function for the continuous phase-type distribution are summarized in Table 1. For the mathematical derivations of these formulas we refer to Bladt and Nielsen (2017).

**Table 1:**
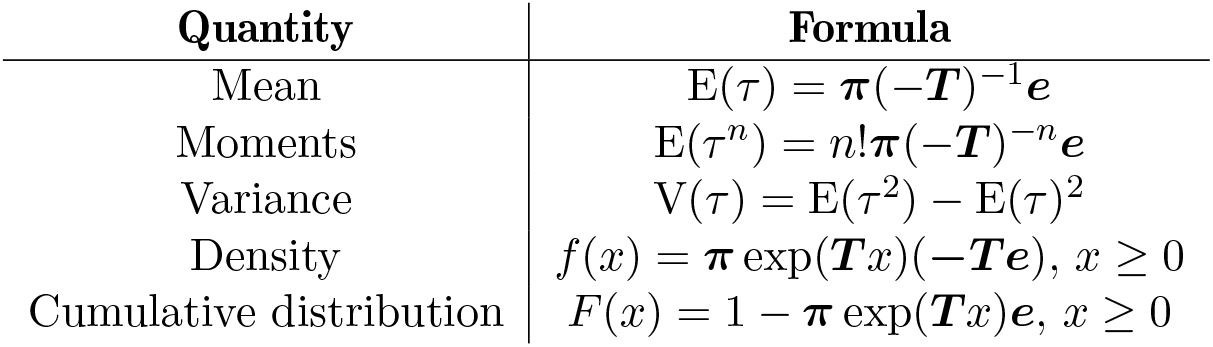
Formulas for the mean, moments, variance, probability density function and cumulative distribution function of the continuous phase-type distribution. Here, ***π*** is the vector of initial probabilities, ***T*** is the subintensity matrix and ***e*** is a vector of ones.

| Quantity | Formula |
| --- | --- |
| Mean | $E(\tau) = \boldsymbol{\pi}(-\mathbf{T})^{-1}\mathbf{e}$ |
| Moments | $E(\tau^n) = n!\boldsymbol{\pi}(-\mathbf{T})^{-n}\mathbf{e}$ |
| Variance | $V(\tau) = E(\tau^2) - E(\tau)^2$ |
| Density | $f(x) = \boldsymbol{\pi} \exp(\mathbf{T}x)(-\mathbf{T}\mathbf{e}), x \geq 0$ |
| Cumulative distribution | $F(x) = 1 - \boldsymbol{\pi} \exp(\mathbf{T}x)\mathbf{e}, x \geq 0$ |

Continuous phase-type distributions can be linearly transformed via rewards (Bladt and Nielsen 2017). This is achieved by assigning a non-negative reward to each of the transient states 1, …, *p*. The resulting distribution is also a phase-type distribution. Let the rewards be given by the function *r*(*i*), *i* = 1, …, *p*, and summarized in the vector ***r*** = (*r*_1_, …, *r*_*p*_), where *r*(*i*) = *r*_*i*_. Consider the reward-transformed random variable

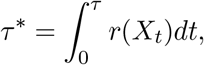

where {*X*_*t*_}_*t*≥0_ is the underlying jump Markov process for the original phase-type distribution *τ* ∼ PH(***π, T***). We have that *τ* ^*^ is phase-type distributed with ***π***^*^ and ***T*** ^*^ denoting the initial distribution and subintensity matrix of this distribution, respectively. If all the elements in the reward vector ***r*** are strictly positive, then ***T*** ^*^ = diag(**1*/r***)***T***, where diag(**1*/r***) is the *p* × *p* diagonal matrix with 1*/r*_*i*_ on the diagonal, while the initial probability vector stays the same (***π***^*^ = ***π***). If ***r*** contains zero-valued rewards, then the states should with a reward of zero can be excluded. As a result, the transient states should be re-defined and the resulting ***π***^*^ and ***T*** ^*^ are lower-dimensional; we refer to Theorem 3.1.33 in Bladt and Nielsen (2017) for the mathematical details.

If several univariate continuous phase-type distributions are defined by the same subintensity matrix but different reward vectors (***r***_**1**_, ***r***_**2**_, …, ***r***_***m***_), then we represent the system as a multivariate continuous phase-type distribution MPH(***π, T***, ***R***), where ***π*** is the initial probability vector, ***T*** is the subintensity matrix, and ***R*** = (***r***_**1**_, ***r***_**2**_, …, ***r***_*m*_) is the *p* × *m* reward matrix.

### 2.2. Defining the phase-type object

To exemplify the usage of **PhaseTypeR**, we will use phase-type representations of common summary statistics in population genetics. Perhaps the most prominent example is the time until the most recent common ancestor *T*_MRCA_, which can be defined as a convolution of exponential distributions following the standard coalescent process (see Figure 1). This means that the *T*_MRCA_ follows a univariate continuous phase-type distribution *T*_MRCA_ ∼ PH(***π, T***), with initial probabilities ***π*** and subintensity matrix ***T*** (Hobolth *et al*. 2019). For the standard coalescent model (Kingman 1982) and a sample size of four chromosomes, the subintensity matrix for the *T*_MRCA_ is given by

**Figure 1:**
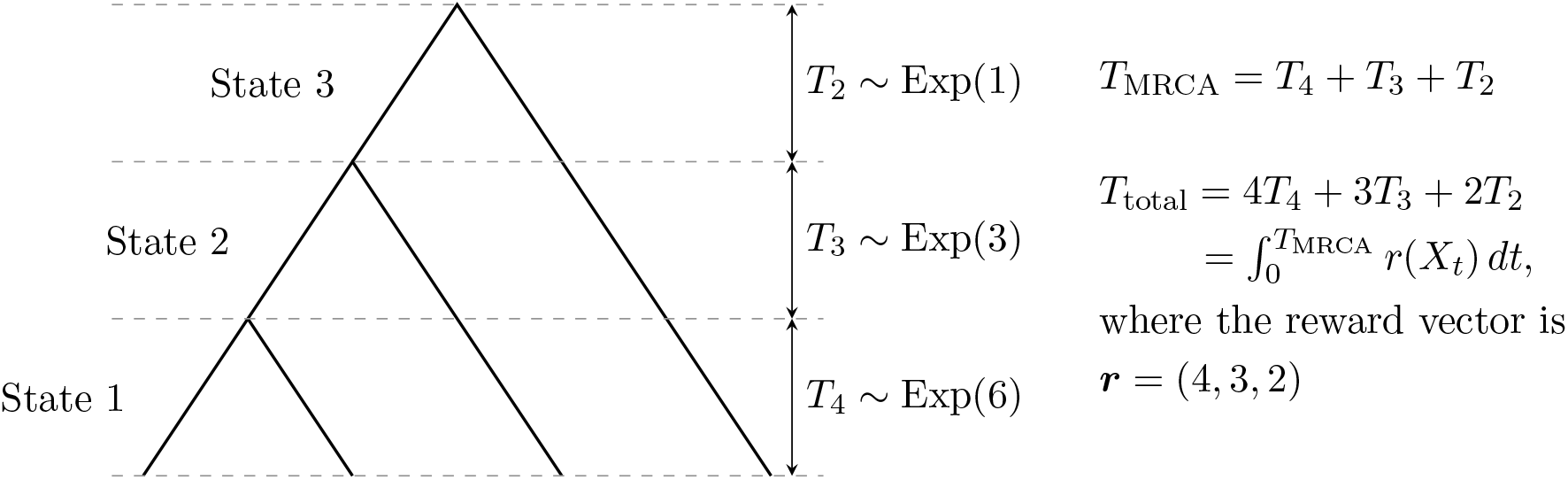
Coalescent process for *n* = 4 samples. The underlying Markov jump process {*X*_*t*_} is in state *X*_*t*_ = 1 for 0 ≤ *t < T*_4_, *X*_*t*_ = 2 for *T*_4_ ≤ *t < T*_4_ + *T*_3_, and *X*_*t*_ = 3 for *T*_4_ + *T*_3_ ≤ *t < T*_4_ + *T*_3_ + *T*_2_. The process is in the absorbing state for *t* ≥ *T*_4_ + *T*_3_ + *T*_2_. The rewards in the states correspond to the number of lineages and the reward-transformed variable *T*_total_ correspond to the total tree length.

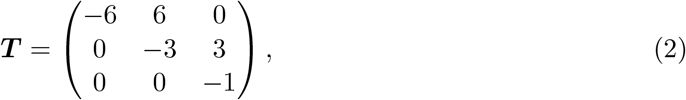

with initial probability vector ***π*** = (1, 0, 0).

We can specify the initial probabilities and the subintensity matrix for this univariate continuous phase-type distribution using the `PH()` function:

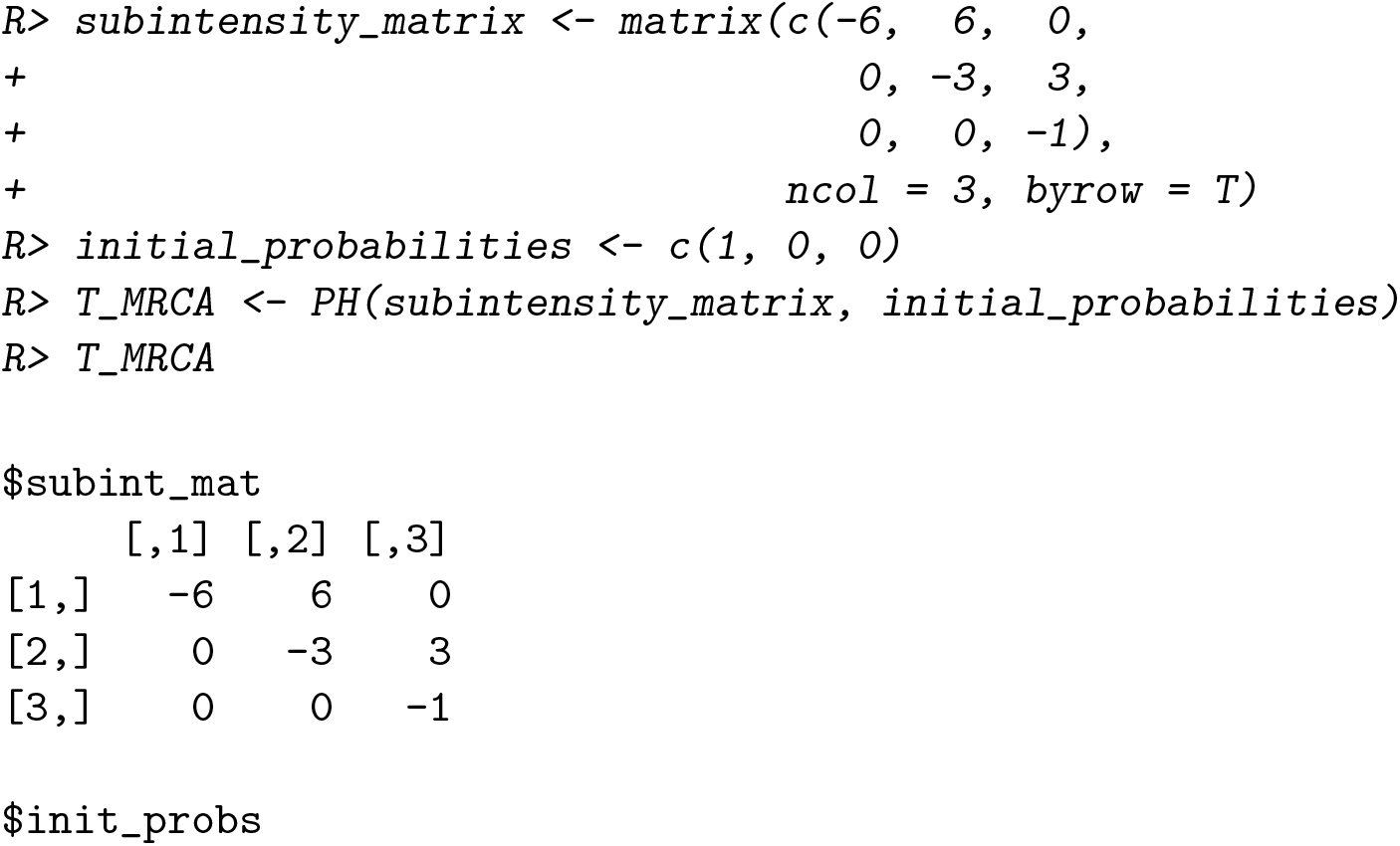

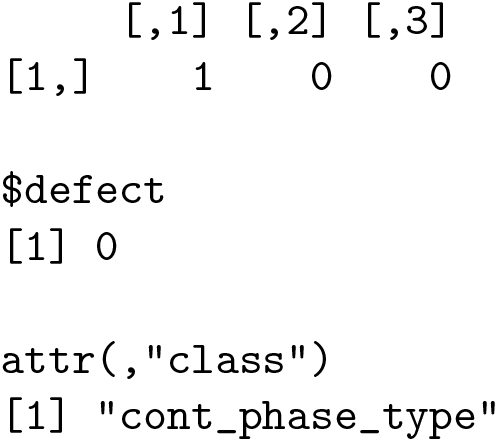

### 2.3. The mean and the variance of a phase-type distribution

The mean and variance of a phase-type object can be accessed by `mean()` and `var()`, respectively. For the phase-type representation of *T*_MRCA_ defined above, `mean(T_MRCA)` yields1.5 and `mean(T_MRCA)` yields 1.138889, which match the well-known results from classical population genetics formulas (Wakeley 2009, Section 3.3).

### 2.4. Distribution functions and random sampling for a phase-type distribution

**PhaseTypeR** uses standard R suffixes for the probability density function (`dPH`), the cumulative distribution function (`pPH`), the quantile function (`qPH`) and the random sampling function (`rPH`) for univariate continuous phase-type distributions:

~~~
*R> dPH(c(0*.*1, 0*.*5, 0*.*8), T_MRCA)*
[1] 0.06482665 0.48210919 0.54651397
*R> pPH(c(0*.*1, 0*.*5, 0*.*8), T_MRCA)*
[1] 0.002348541 0.121417559 0.280279868
*R> qPH(c(0*.*05, 0*.*5, 0*.*95), T_MRCA)*
[1] 0.3302855 1.2328314 3.5830871
*R> set*.*seed(3)
R> rPH(3, T_MRCA)*
[1] 0.6459884 0.1019513 1.0577725
~~~

Sometimes, it is of interest to retrieve the full sample path of the Markov jump process. The user can achieve so using the `rFullPH` function, which returns a data frame containing all the visited states and the time spent in each:

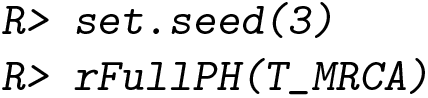

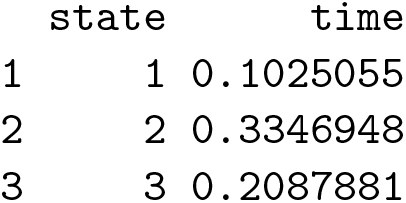

### 2.5. Reward transformation

The *T*_MRCA_ is tightly related to the total tree length, or *T*_total_ (Hobolth *et al*. 2019). More specifically, *T*_total_ is a linear transformation of *T*_MRCA_, so its phase-type representation can be obtained by a reward transformation. The reward vector needed is ***r*** = (*n, n* − 1, …, 2), where *n* is the sample size (recall Figure 1).

Reward transformation in **PhaseTypeR** can be done using the `reward_phase_type()` function. For the case of *n* = 4 the reward vector is ***r*** = (4, 3, 2), and a phase-type representation of *T*_total_ can be obtained by reward-transforming *T*_MRCA_:

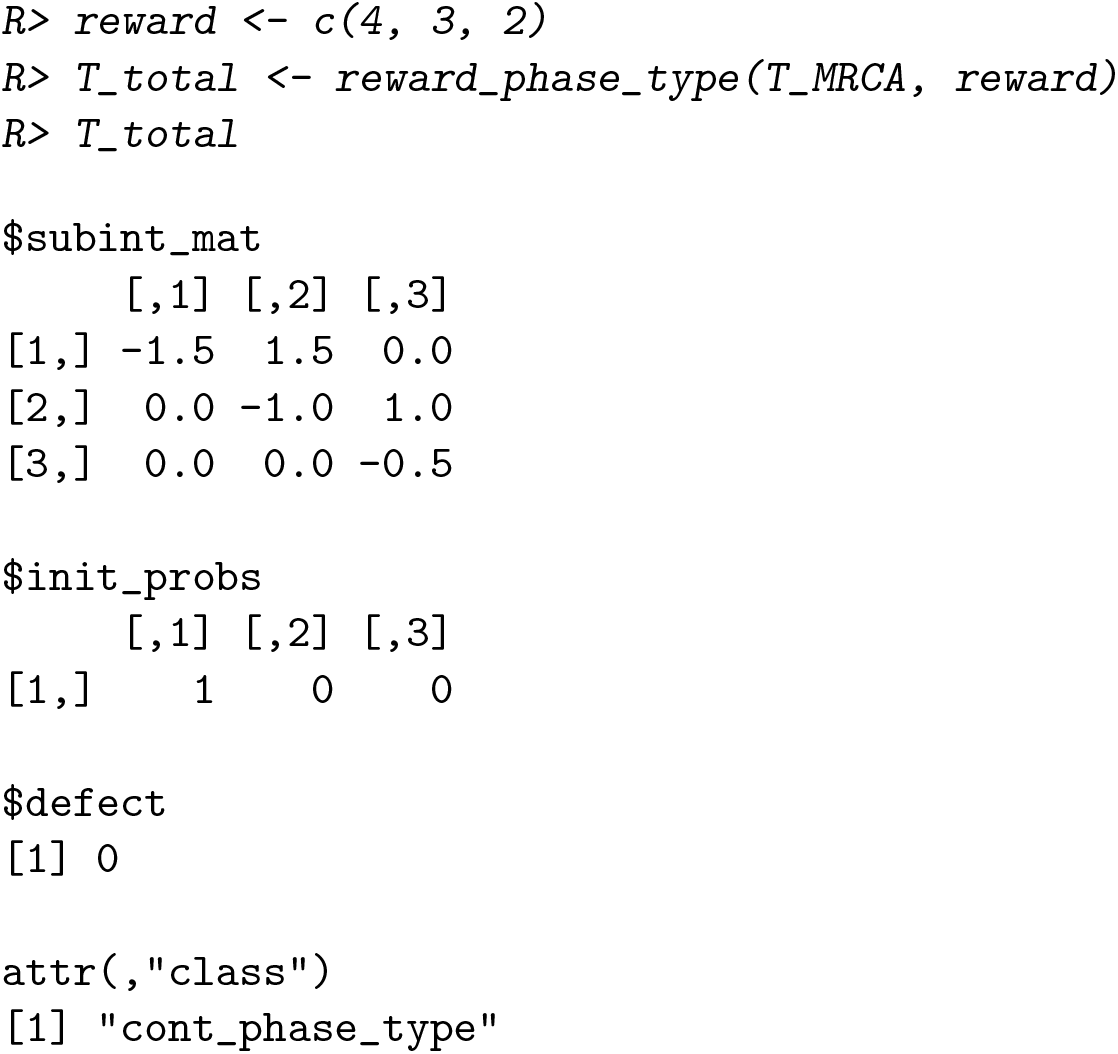

The mean and variance of the total branch length are given by

~~~
*R> c(mean(T_total), var(T_total))*
[1] 3.666667 5.444444
~~~

We note once again that these results match the ones derived from classical population genetic formulas (e.g. Wakeley 2009, Section 3.3).

### 2.6. The multivariate continuous phase-type distribution

Similar to the construction of *T*_total_, we can reward-transform the phase-type representation of *T*_MRCA_ to get the distribution of the total branch length leading to each of the elements of the site frequency spectrum (*ξ*_1_, …, *ξ*_*n*−1_), i.e. singletons (*ξ*_1_), doubletons (*ξ*_2_), etc. In order to do so for *n* = 4, we first need to extend the subintensity matrix ***T*** by sub-dividing state 3 into two. The reason for this is that 1/3 of the times the second coalescent will lead to the creation of two doubleton branches, while 2/3 of the times it will lead to one singleton branch and one tripleton branch (see Figure 2). The resulting subintensity matrix is given by

**Figure 2:**
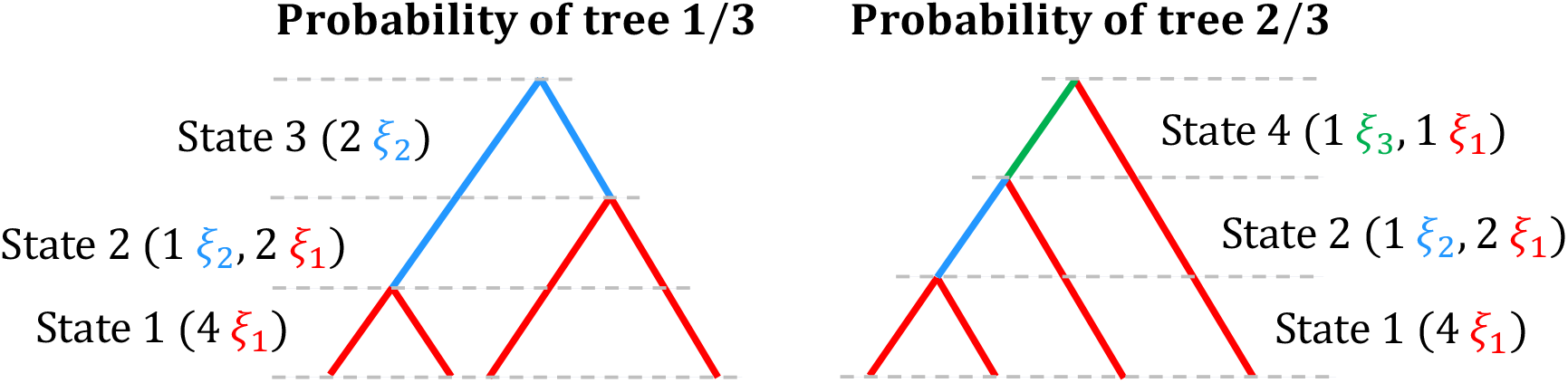
Coalescent process for *n* = 4 samples. State 1 contains 4 singleton branches (mutations in singleton branches are only present in one sample, represented in red), state 2 contains 2 singleton branches and 1 doubleton branch (mutations in doubleton branches are present in two samples, represented in blue), state 3 contains 2 doubleton branches, and state 4 contains 1 singleton and 1 tripleton branch (mutations in tripleton branches are present in three samples, represented in green).

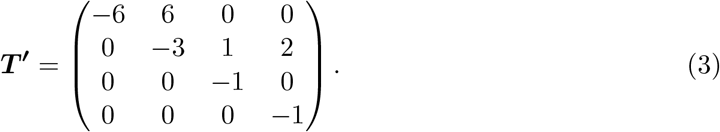

Note that this is a less efficient but equal phase-type representation of *T*_MRCA_ if the initial probability vector is ***π′***= (1, 0, 0, 0).

Following Hobolth *et al*. (2019), if we transform *T*_MRCA_ with a reward of ***r***_**1**_ = (4, 2, 0, 1), we get a phase-type representation of the total branch length leading to singletons, denoted *L*_1_. Knowing that the reward vectors for doubletons and tripletons are ***r***_**2**_ = (0, 1, 2, 0) and ***r***_**3**_ = (0, 0, 0, 1), instead of reward-transforming each element of the site frequency spectrum separately, we can define a multivariate continuous phase-type distribution ***L*** = (*L*_1_, *L*_2_, *L*_3_) ∼ MPH(***π***^***′***^, ***T***^***′***^, ***R***) with initial distribution ***π′***= (1, 0, 0, 0), subintensity matrix ***T′*** and reward matrix ***R*** = (***r***_**1**_, ***r***_**2**_, ***r***_**3**_).

Multivariate continuous phase-type distributions are implemented in **PhaseTypeR** as follows:

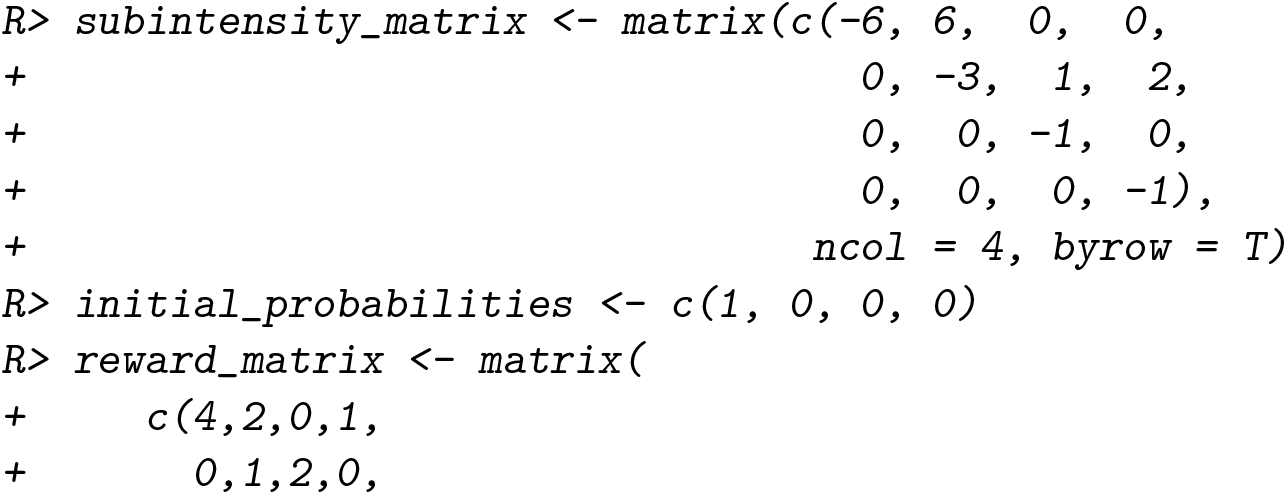

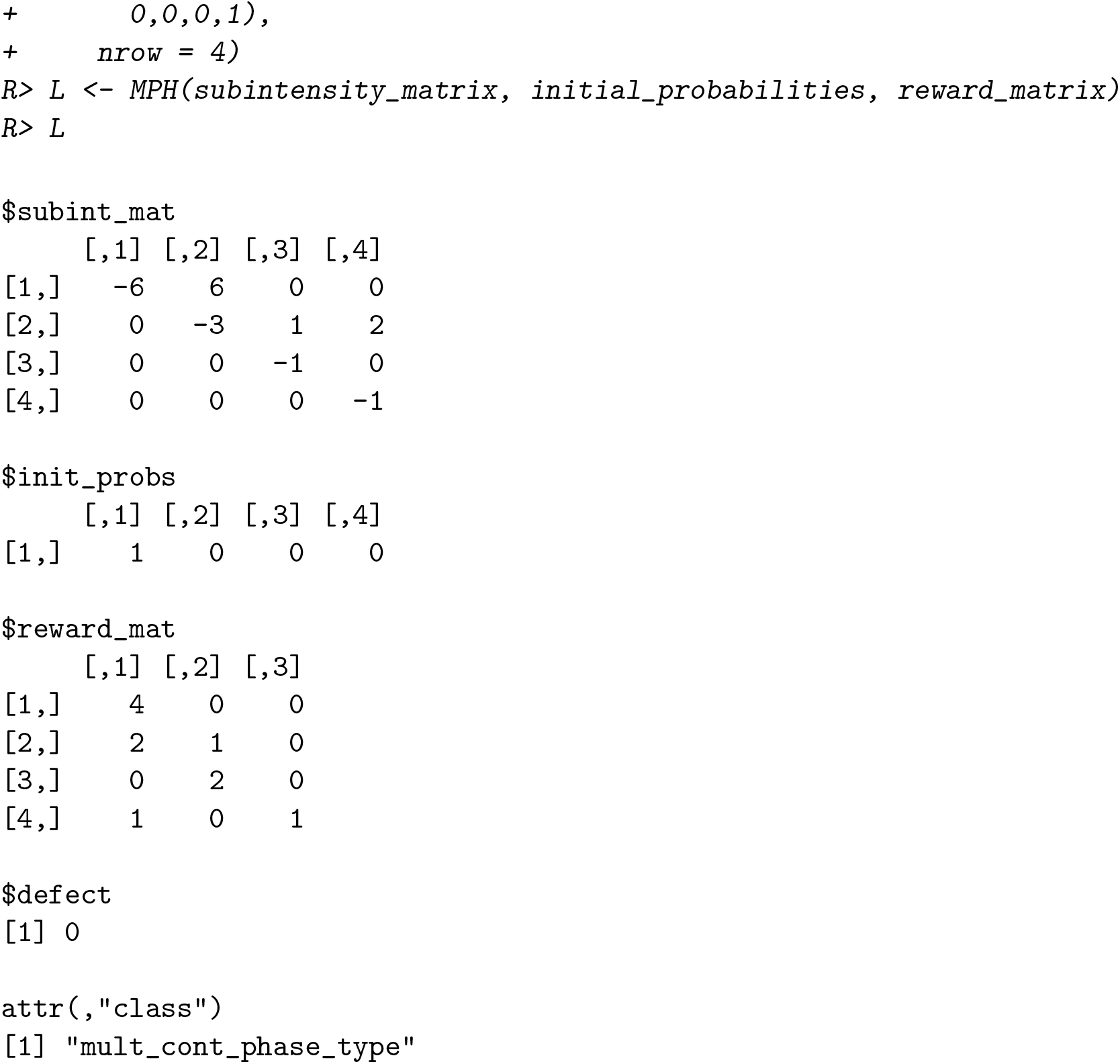

This type of multivariate representation is useful to calculate the variance-covariance matrix of the variables. For the multivariate case, this can be accessed using `var()`:

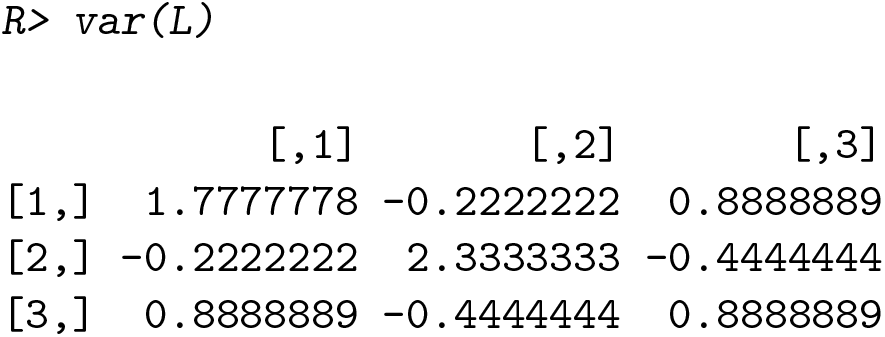

Here, the diagonal is the variance of each of the elements (total branch length leading to singletons, doubletons and tripletons in this case), and the off-diagonal values are the covariances between the different elements. The analytical formula for calculating the covariance can be found in Theorem 8.1.5 in Bladt and Nielsen (2017).

Moreover, **PhaseTypeR** also computes univariate quantities related to the marginal distributions of the MPH, i.e. the probability density function (`dMPH`), the cumulative distribution function (`pMPH`) and the quantile function (`qMPH`). Random draws (`rMPH`) and random draws with full path (`rFullMPH`) for the multivariate case use the same underlying sample path for the Markov jump process.

## 3. Discrete phase-type distributions

### 3.1. Theory for the discrete phase-type distribution

A discrete phase-type distribution describes a process of geometric distributions that occur sequentially until absorption. It is similar to the continuous case, but the underlying process is a discrete time absorbing Markov chain instead of a Markov jump process.

If we define the Markov chain as {*X*_*n*_}_*n*∈ℕ_, which has *p* transient states and one absorbing state, the discrete time until absorption *τ* follows a discrete phase-type distribution. The transition probability matrix ***P*** of the Markov chain is then defined as

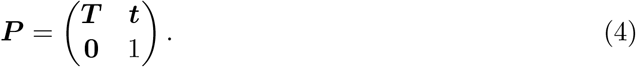

The transition probabilities among the transient states are therefore contained in the *p* × *p* subtransition matrix ***T***. Similar to the continuous case, the discrete phase-type distribution can be described solely by ***T***. Since all the rows in ***P*** sum to 1 we have ***P e*** = ***e*** and the exit probability vector is given by ***t*** = ***e*** − ***T e*** = (***I*** − ***T***)***e***, where ***I*** is a *p* × *p* identity matrix. If we let ***π*** be the vector of initial probabilities, then *τ* ∼ DPH(***π, T***).

Similar to the continuous case, the mean, variance, probability density function and cumulative distribution function for the discrete phase-type distribution can be defined using matrix manipulation (Bladt and Nielsen 2017), and they are summarized in Table 2.

**Table 2:** Formulas for the mean, moments, variance, probability density and cumulative distribution function of the discrete phase-type distribution. Here, ***π*** is the vector of initial probability, ***T*** is the subtransition matrix, ***t*** is the exit probability vector, ***e*** is a vector of ones, and ***I*** is an identity matrix.

| Quantity | Formula |
| --- | --- |
| Mean | $\text{E}(\tau) = \boldsymbol{\pi}(\mathbf{I} - \mathbf{T})^{-1}\mathbf{e}$ |
| Moments | $\text{E}(\tau^n) = n! \boldsymbol{\pi}(\mathbf{I} - \mathbf{T})^{-n}\mathbf{e}$ |
| Variance | $\text{V}(\tau) = \text{E}(\tau^2) - \text{E}(\tau)^2$ |
| Density | $f(x) = \boldsymbol{\pi}\mathbf{T}^{x-1}\mathbf{t}, x \geq 1$ |
| Cumulative distribution | $F(x) = 1 - \boldsymbol{\pi}\mathbf{T}^x\mathbf{e}, x \geq 1$ |

Additionally, discrete phase-type distributions can also be transformed with non-negative integer rewards. If a reward for a certain state is set to 0, then that state is removed from the subtransition matrix. If instead it is set to a positive integer larger than 1, then the subtransition matrix is extended to “force” the Markov chain to pass through a state several times. The full mathematical construction of reward transformations for the discrete case is presented in Theorem 5.2 in Campillo Navarro (2018).

Similar to the continuous case, multivariate discrete phase-type distributions can be constructed by combining univariate distributions that share the same subtransition matrix but have different rewards (***r***_**1**_, ***r***_**2**_, …, ***r***_***m***_). The resulting joint distribution MDPH(***π, T***, ***R***) contains an initial probability vector ***π***, a subtransition matrix ***T*** and a *p* × *m* reward matrix ***R*** = (***r***_**1**_, ***r***_**2**_, …, ***r***_***m***_).

### 3.2. Defining the discrete phase-type object

Another summary statistic in population genetics that can be represented using phase-type theory is the number of segregating sites *S*_total_ (Hobolth *et al*. 2019). Consider the coalescent process with a sample size of *n* = 3, i.e. we have a state 1 with three singleton branches and state 2 with one singleton branch and one doubleton branch (Kingman 1982). The coalescent rate in state 1 is 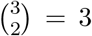 and the mutation rate is *θ/*2 on each of the three branches (so 3*θ/*2 in total). The probability that the first event in the ancestral process is a mutation before a coalescent is given by the mutation rate relative to the total rate, a situation which corresponds to two competing exponential distributions (see e.g. Wakeley 2009, equations (2.60) and (4.5)). Therefore, the number of mutations when three branches are present is geometrically distributed with probability of mutation

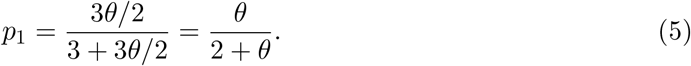

Similarly, when two branches are present, the coalescent rate is 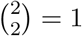 and the total mutation rate is 2*θ/*2. Thus, the number of mutations in this case is geometrically distributed with probability of mutation

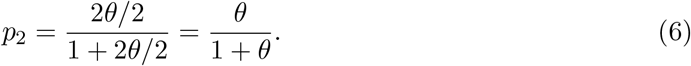

We may describe the situation using a discrete phase-type distribution with subtransition probability matrix

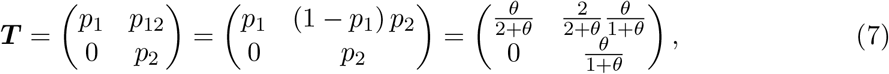

Here, a jump into state 1 corresponds to a mutation on the level of the tree with three branches, and a jump into state 2 corresponds to a mutation on the level of the tree with two branches. We can also start in a situation with no mutation, which corresponds to directly jumping to the absorbing state. In order to model this, we can work with a defective initial distribution given by ***π*** = (*p*_1_, *p*_12_). The probability of zero jumps (mutations) is then

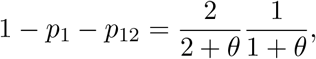

which corresponds to the defect. The total number of mutations, thus, follows a univariate discrete phase-type distribution *S*_total_ ∼ DPH(***π, T***). We remark that this same distribution arises from adding Poisson mutations on the phase-type distributed total tree length (Theorem 3.5, eq. (19) in Hobolth *et al*. (2019)).

In **PhaseTypeR**, it is straightforward to specify a univariate discrete phase-type distributions with `DPH()`. For the case of *S*_total_ when *n* = 3 and *θ* = 3:

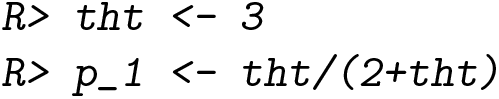

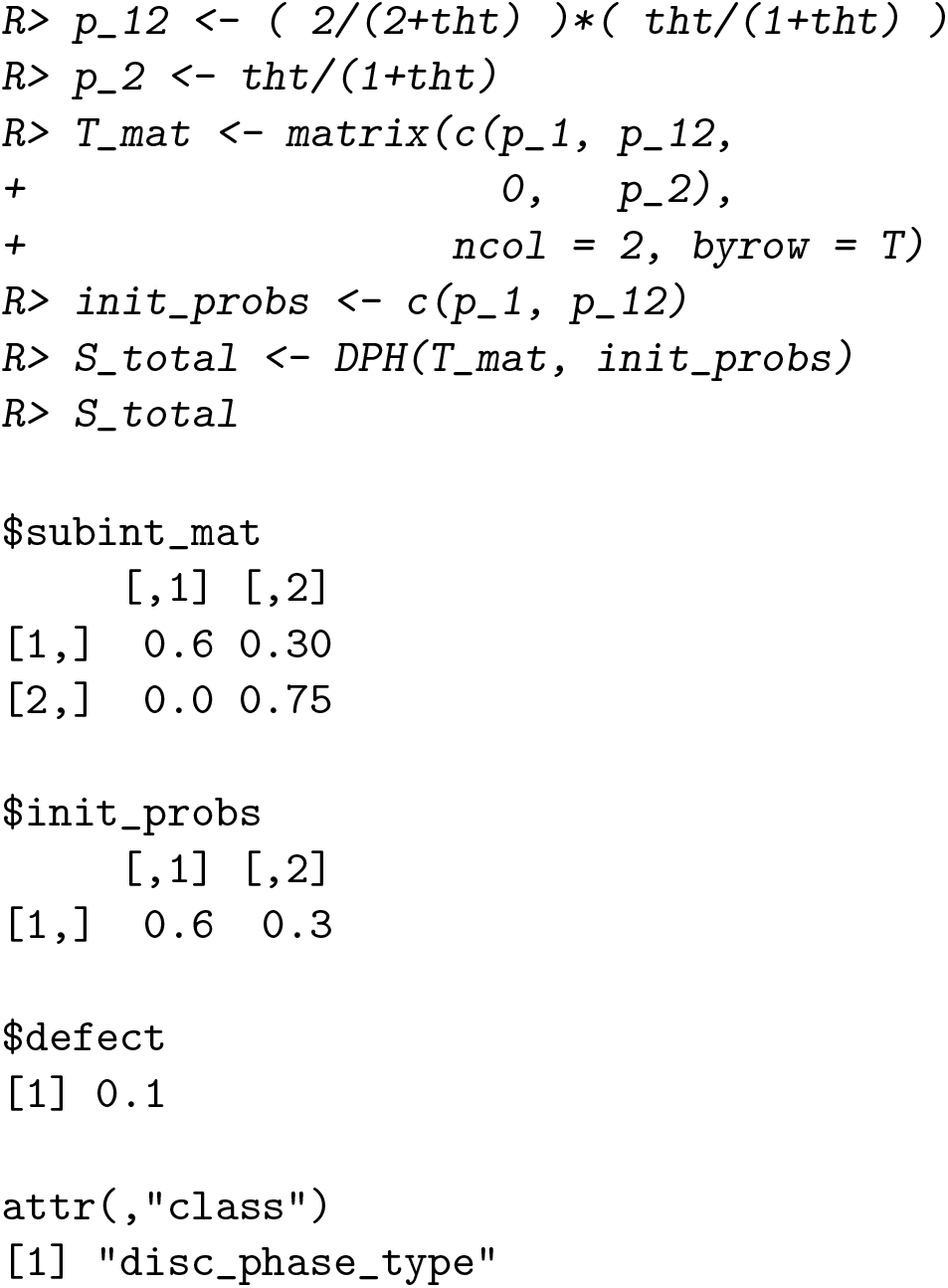

### 3.3. The mean and the variance of a discrete phase-type distribution

Similar to the continuous case, the mean and variance of the discrete phase-type object can be computed by `mean()` and `var()`, respectively. For the phase-type representation of *S*_total_ defined above, `mean(S_total)` yields 4.5 and `var(S_total)` yields 15.75. These results match the ones derived from classical population genetic formulas (e.g. Wakeley 2009, Section 4.1.1).

### 3.4. Distribution functions and sampling for a discrete phase-type distribution

**PhaseTypeR** also contains functions for the probability density function (`dDPH`), the cumulative distribution function (`pDPH`), the quantile function (`qDPH`) and random sampling (`rDPH`) of univariate discrete phase-type distributions:

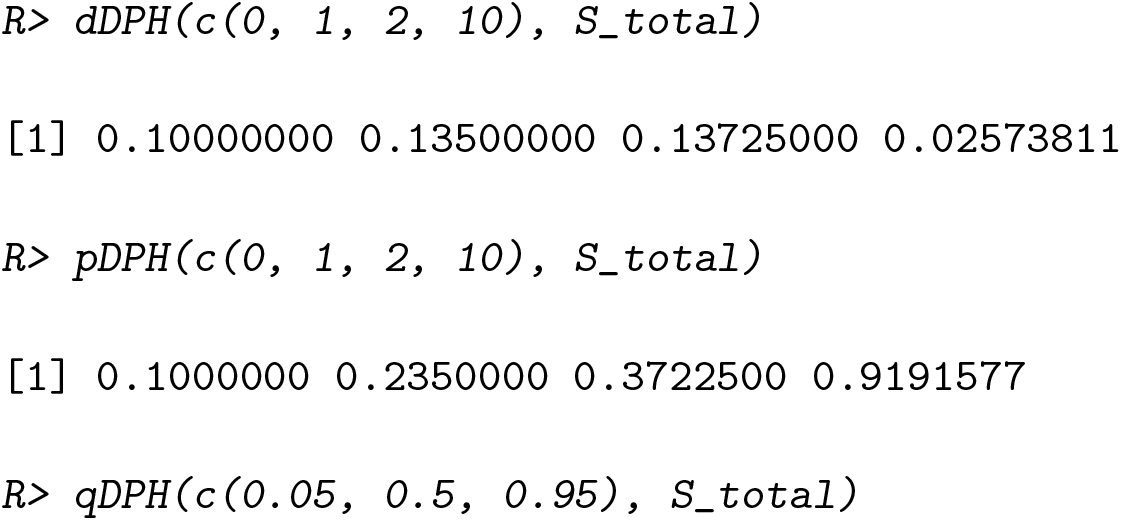

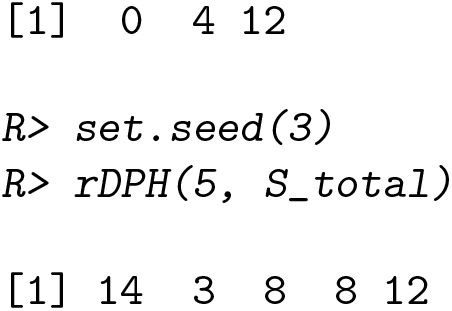

We can also simulate a sample path from the Markov chain by using the `rFullDPH()` function. This returns a data frame with the visited states and the time spent in each of them:

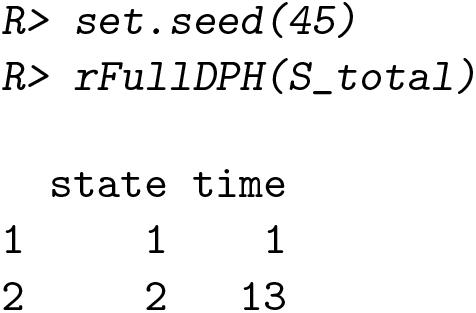

### 3.5. Reward transformation

While *S*_total_ does not distinguish the different types of segregating sites, sometimes we are interested in knowing the type of mutations based on the site frequency spectrum. When *n* = 3, there are two types of segregating sites, namely singletons (*ξ*_1_) and doubletons (*ξ*_2_). Mutations in state 1 are always singletons, but mutations in state 2 are singletons with probability 1*/*2 and doubletons with probability 1*/*2. We therefore extend the subtransition probability matrix to

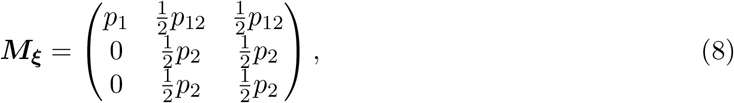

where the new state 2 corresponds to singletons in the old state 2 and the new state 3 corresponds to doubletons in the old state 2. If we define an initial probability vector of 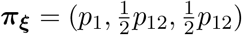, we can define a discrete phase-type distribution *S*_total_ ∼ DPH(***π***_***ξ***_, ***M***_***ξ***_). This way of defining *S*_total_ is a more inefficient though equivalent representation compared to the definition in the previous section. However, we can now transform the phase-type distribution via rewards to get discrete phase-type representations of the different elements of the site frequency spectrum. For example, *ξ*_1_ ∼ DPH(***π***_**1**_, ***M***_**1**_), which can be derived by reward-transforming *S*_total_ with a reward vector of ***r***_**1**_ = (1, 1, 0). This can be done in **PhaseTypeR** using the `reward_phase_type()` function:

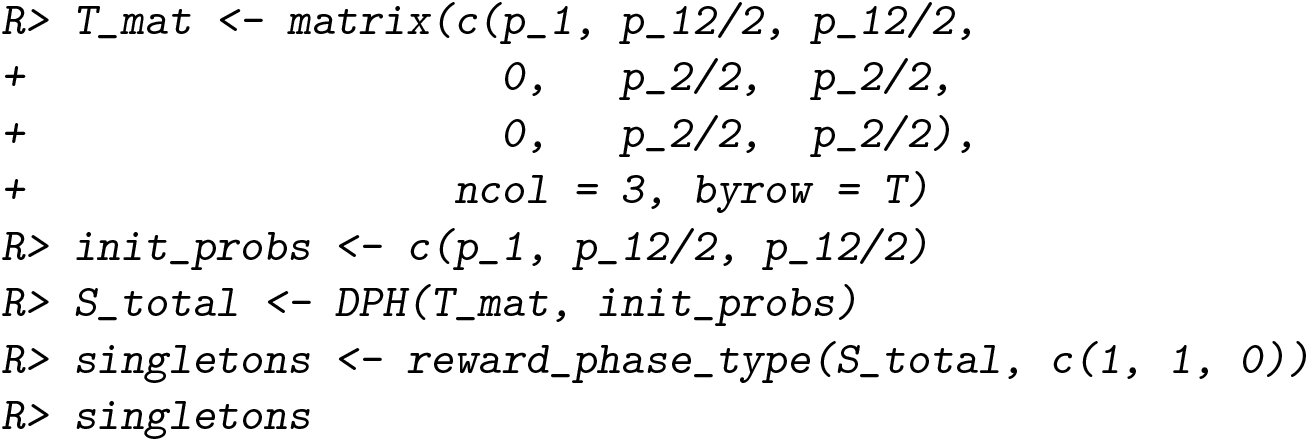

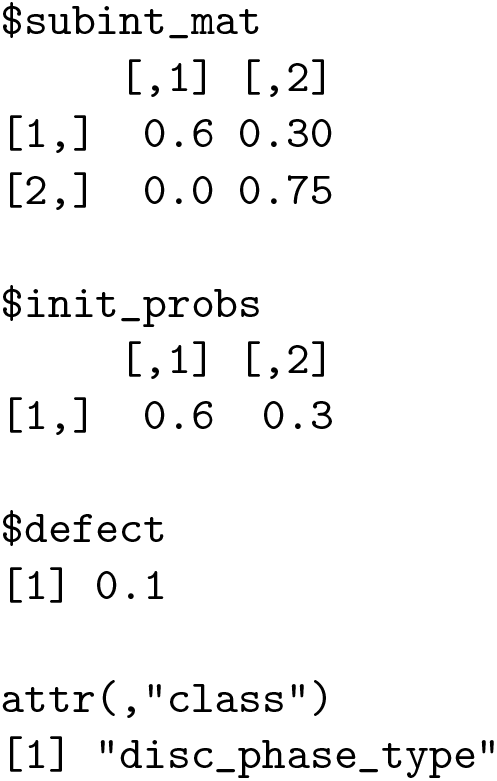

And similarly, for doubletons with a reward vector of ***r***_**2**_ = (0, 0, 1):

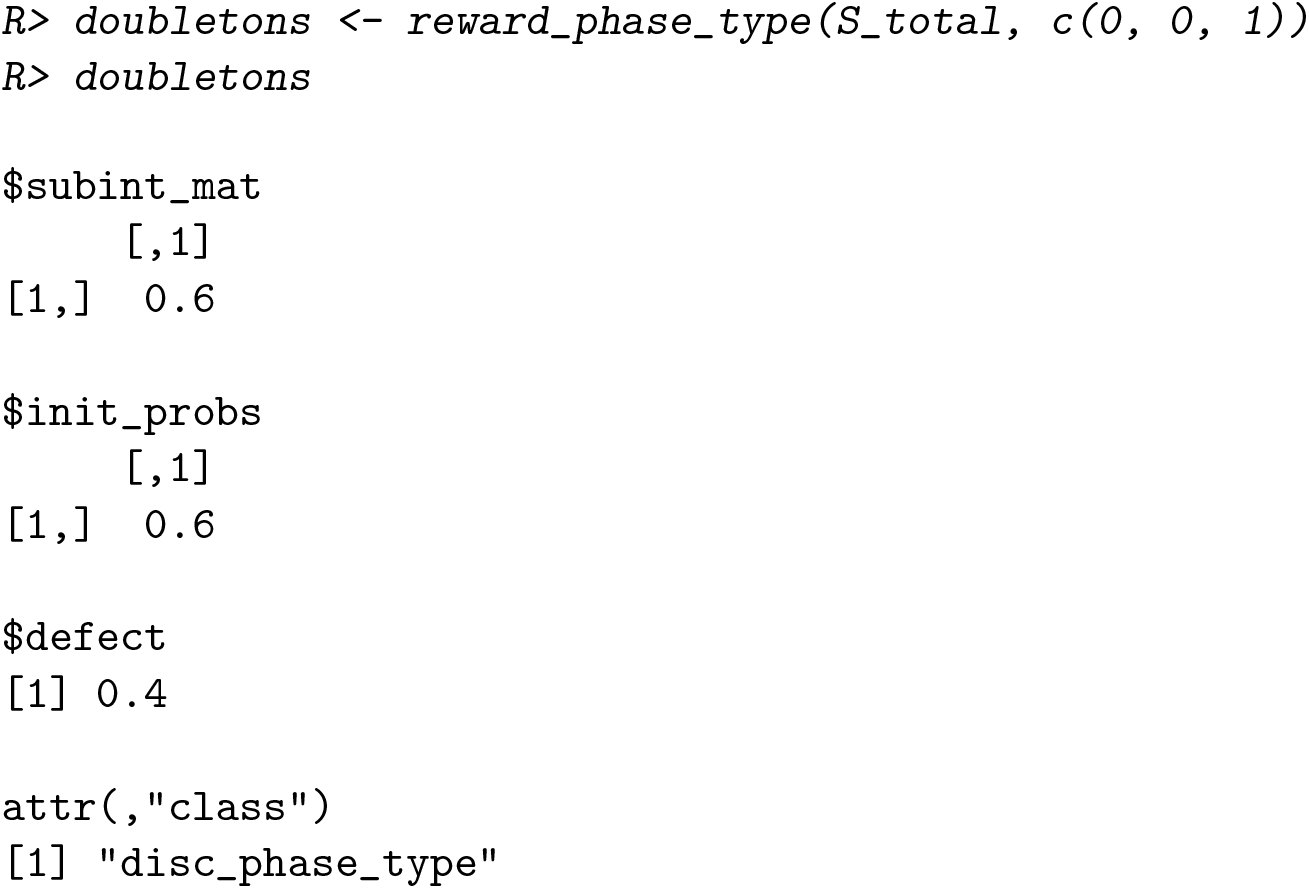

Note that when *θ* = 3

~~~
*R> c(mean(singletons), mean(doubletons))*
[1] 3.0 1.5
~~~

This matches the famous result from coalescent theory (e.g. Wakeley 2009, Section 4.1.3), which states that the mean of the elements in the site frequency spectrum is E[*ξ*_*i*_] = *θ/i, i* = 1, …, *n* − 1.

### 3.6. The multivariate discrete phase-type distribution

Naturally, the joint site frequency spectrum (*ξ*_1_, *ξ*_2_) is multivariate discrete phase-type distributed with initial distribution ***π*** = (*p*_1_, *p*_12_), subtransition matrix ***M***_***ξ***_, and reward vectors ***r***_**1**_ = (1, 1, 0) and ***r***_**2**_ = (0, 0, 1), i.e.

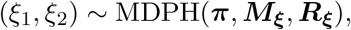

with ***R***_***ξ***_ = (***r***_**1**_, ***r***_**2**_).

Using **PhaseTypeR**:

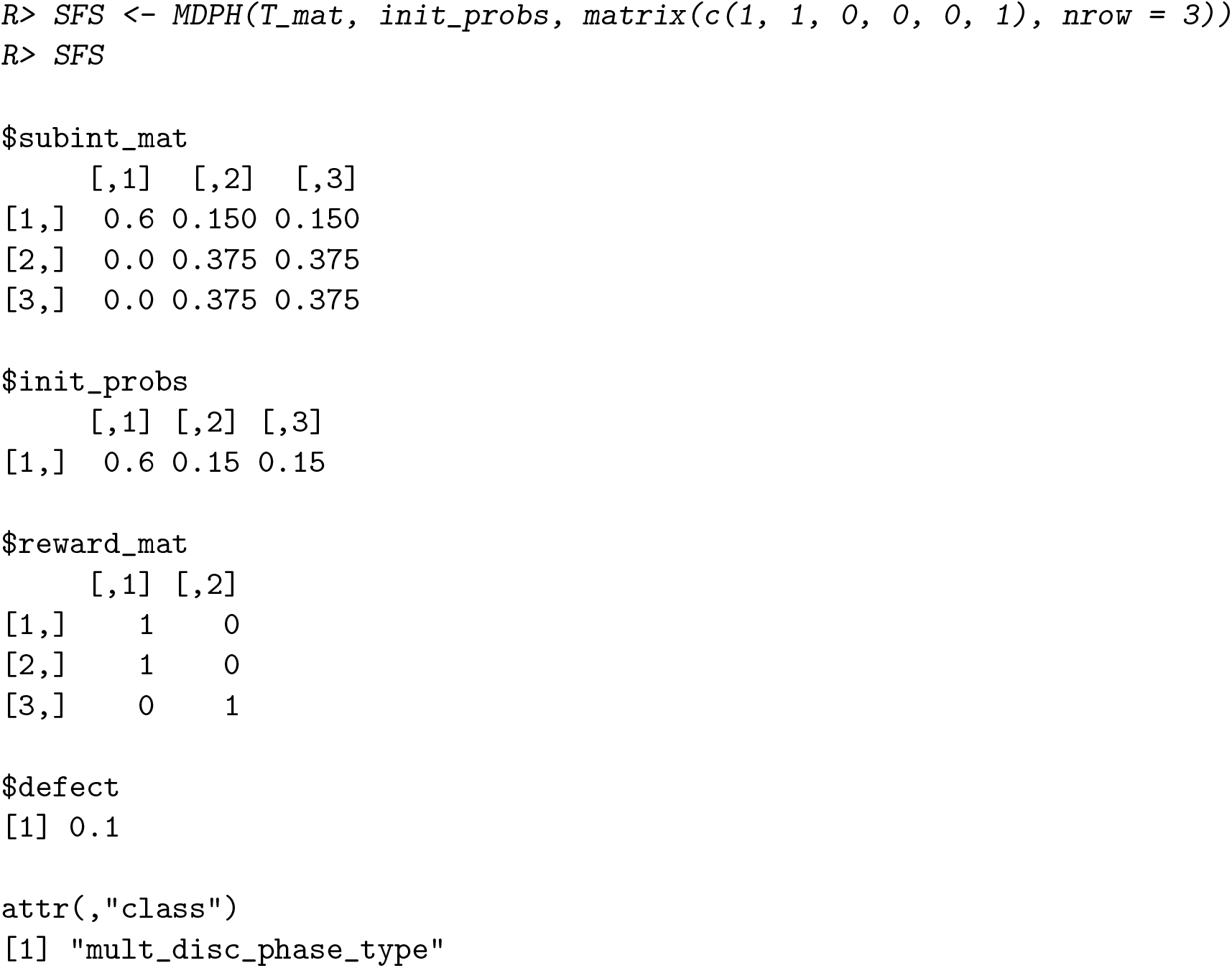

This construction can be extended to any sample size *n* with site frequency spectrum (SFS) (*ξ*_1_, …, *ξ*_*n*−1_). The general situation is described in Hobolth *et al*. (2021), and the special case *n* = 4 is illustrated in detail in Section 4 in that paper.

**PhaseTypeR** can be used to calculate the variance-covariance matrix of a multivariate discrete phase-type distribution:

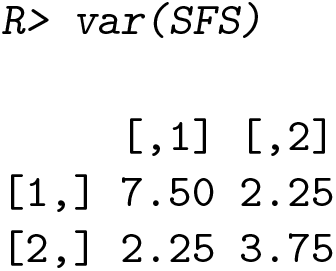

Here, the diagonal corresponds to the variance in the number of singletons and doubletons, respectively. The covariance is provided in the off-diagonal values, whose formula can be consulted in Campillo Navarro (2018).

Similar to the continuous case, **PhaseTypeR** also contains functions for calculating univariate quantities of the marginal distributions of MDPH. These include the probability density function (`dMDPH`), the cumulative distribution function (`pMDPH`) and the quantile function (`qMDPH`). Moreover, random draws (`rMDPH`) and random draws with full path (`rFullMDPH`) for the multivariate discrete case use the same underlying sample path for the Markov chain.

## 4. The coalescent with recombination

The traditional procedure for deriving the correlation between the branch lengths in two loci for a sample of size two is by a first-step analysis (e.g. Wakeley 2009, Section 7). In this section we demonstrate how to use phase-type theory to obtain the result.

The state space and transition rates for the two-locus ancestral recombination graph is shown in Figure 3. The filled circles represent material ancestral to the sample, and the crosses indicate that the most common ancestor has been found. The lines between the circles or crosses indicate if the ancestral material is present on the same chromosome. The starting state is state 1 at present day with two samples from the same chromosome.

**Figure 3:**
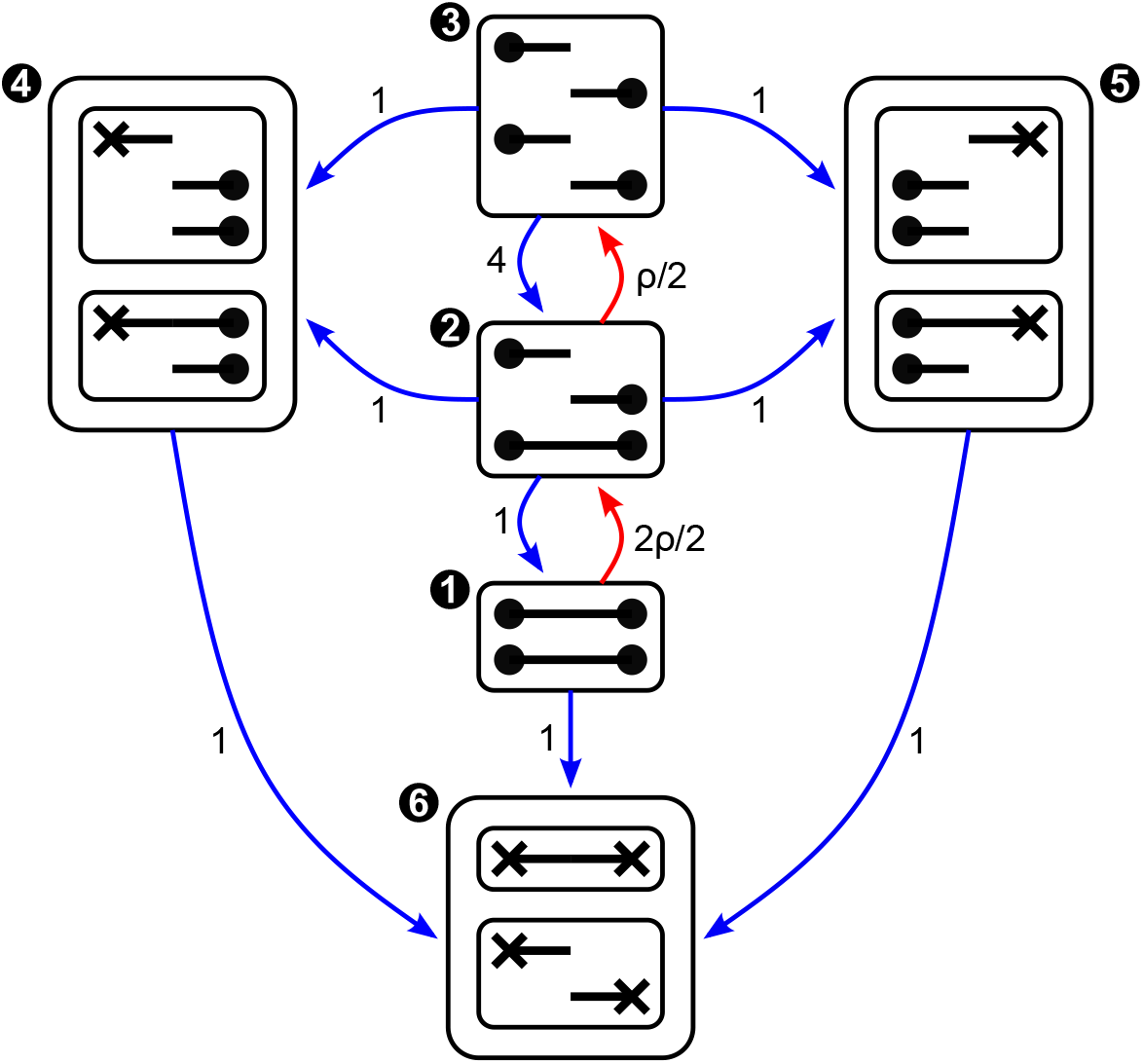
State space and transition rates for the two-locus ancestral recombination graph (ARG). Coalescent events are represented with blue arrows, while recombination events are marked in red. The corresponding coalescent or recombination rates are labeled next to each arrow.

The time *τ* when both loci have found their common ancestor is PH(***α, S***) distributed with ***α*** = (1, 0, 0, 0, 0) and

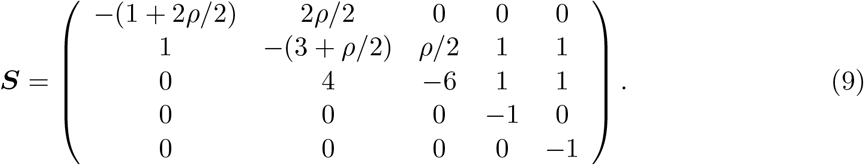

The tree height *T*_left_ in the left locus is the first time the ancestral process {*X*(*t*) : *t* ≥ 0} enters state 4 or state 6 or, equivalently, the time spent in state 1, 2, 3 and 5 before absorption in state 6. We therefore have

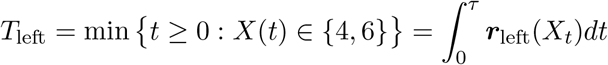

with the reward vector ***r***_left_ = (1, 1, 1, 0, 1). Similarly, the tree height *T*_right_ in the right locus is the first time the ancestral process enters state 5 or state 6 or, equivalently, the time spent in state 1, 2, 3 and 4 before absorption in state 6. We therefore have

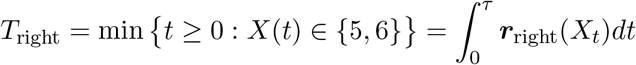

with the reward vector ***r***_right_ = (1, 1, 1, 1, 0). A classical result in population genetics gives the covariance between the two tree heights

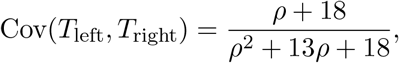

and we note that for large recombination rates Cov(*T*_left_, *T*_right_) is close to zero, and for small recombination rates it is close to one. Note that *T*_left_ and *T*_right_ are both exponentially distributed with a rate of 1, so Var(*T*_left_) = Var(*T*_right_) = 1, and, consequently, Cor(*T*_left_, *T*_right_) = Cov(*T*_left_, *T*_right_) (see also Wakeley 2009, equation (3.10)). Moreover, as shown by a simple proof in Wilton, Carmi, and Hobolth (2015), we have that *P* (*T*_left_ = *T*_right_) = Cov(*T*_left_, *T*_right_).

An implementation using **PhaseTypeR** simply consists of specifying the initial distribution, rate matrix for the ancestral process, rewards for the two tree heights, and calling the variance function for the multivariate phase-type distribution.

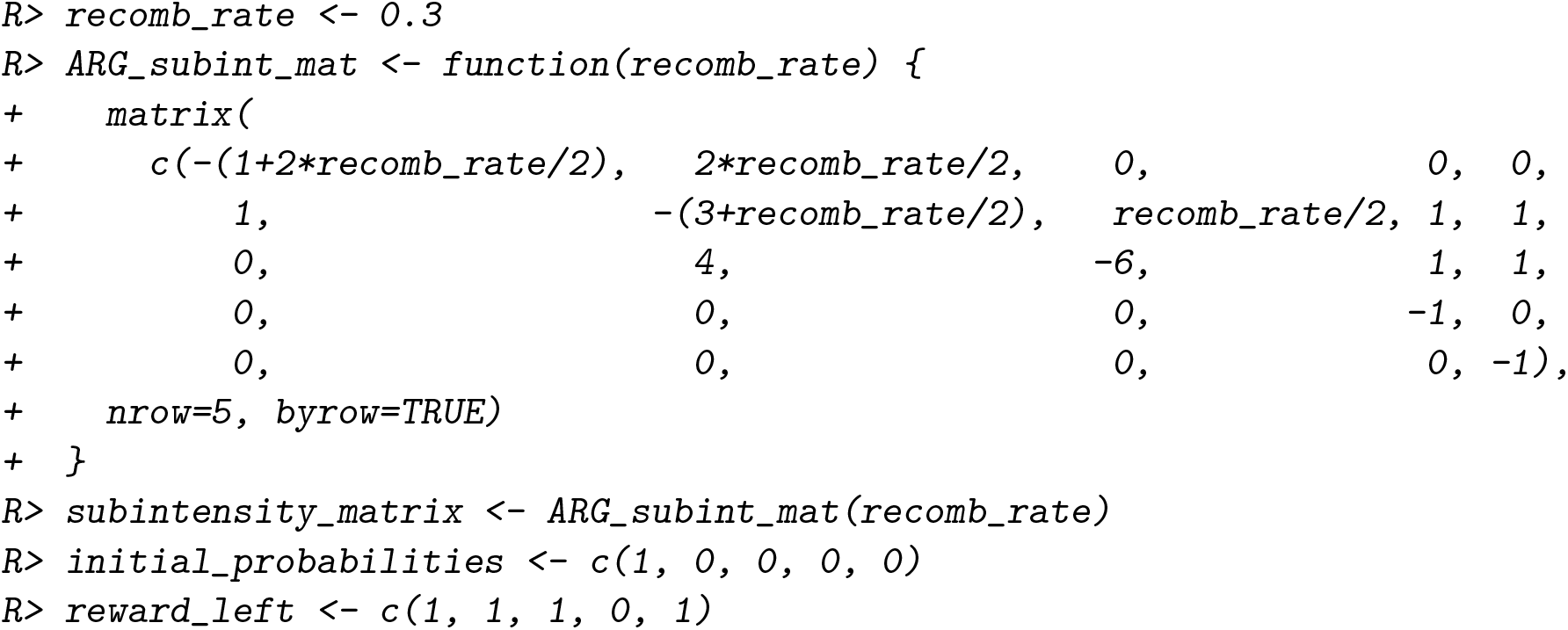

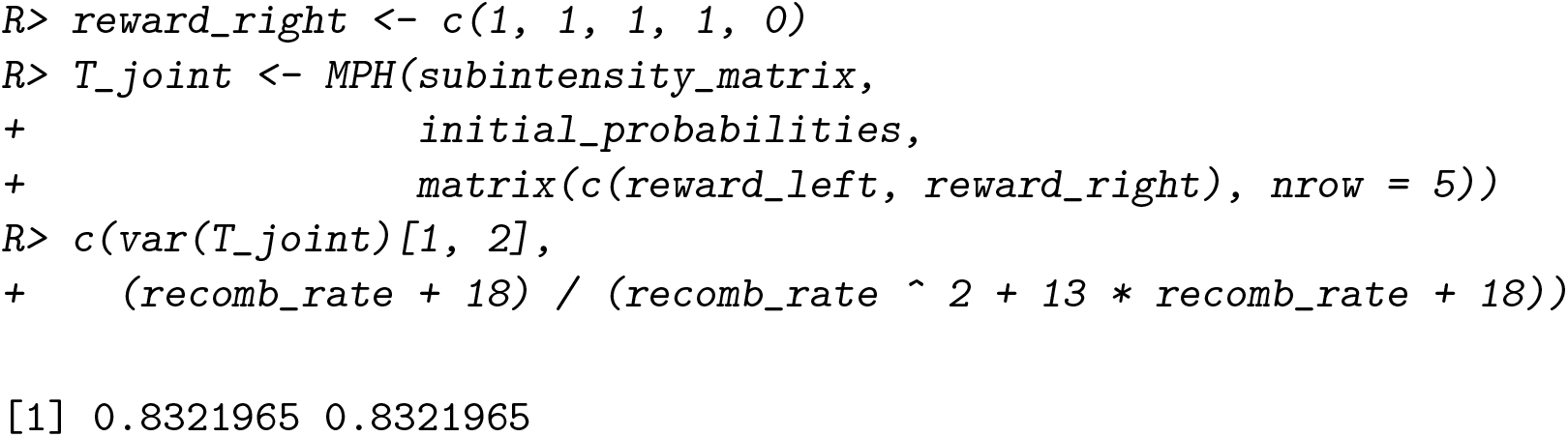

We can see that the phase-type result is equal to the classical formula provided above. From this multivariate phase-type representation of the ARG, we can simulate, for example, 1,000 draws from the joint distribution of (*T*_left_, *T*_right_) using `rMPH(1000, T_joint)` in **PhaseTypeR**. If the recombination rate *ρ* is set to a small value, then most of the draws will result in *T*_left_ = *T*_right_, and the joint density will concentrate along the diagonal, as shown in Figure 4, left (Simonsen and Churchill 1997). If instead *ρ* is large, then most of the draws will result in *T*_left_ ≠ *T*_right_ (Figure 4, right).

**Figure 4:**
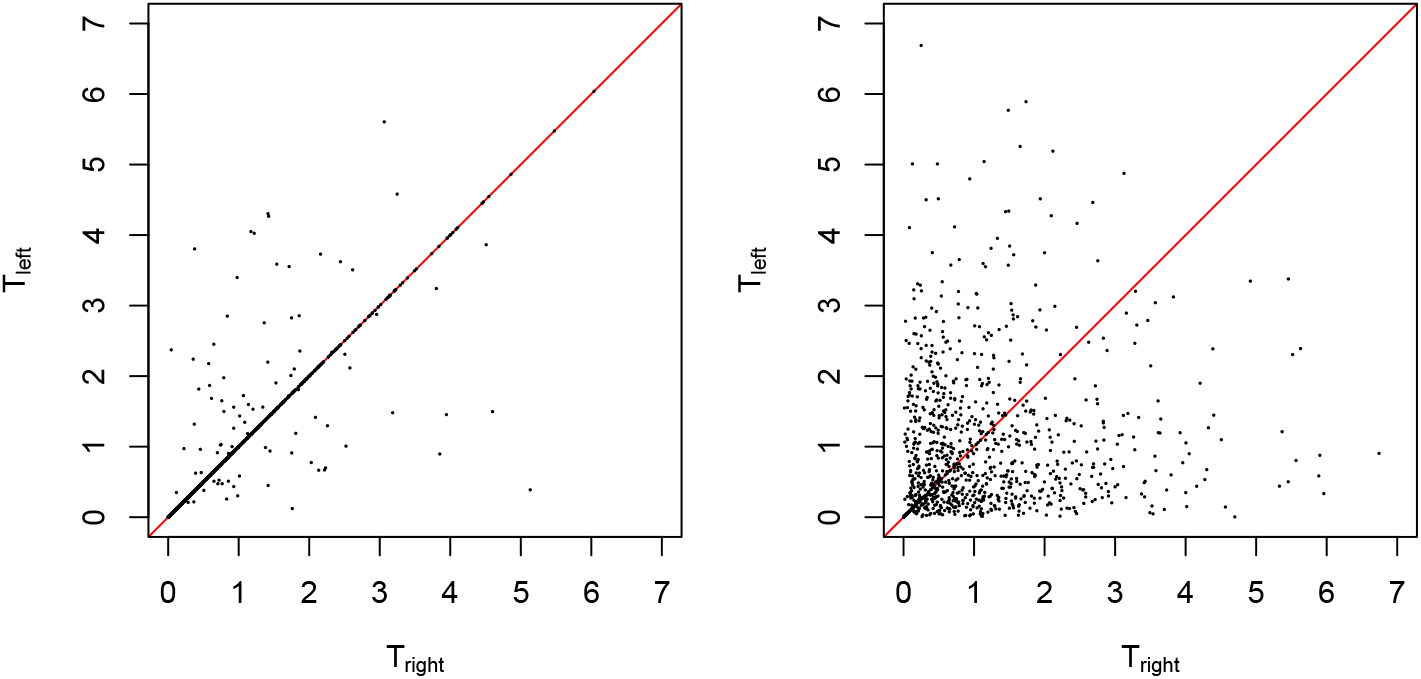
Scatter plot of a simulation of 1,000 draws from the joint distribution of the coalescent times (*T*_left_, *T*_right_) in the two-loci, two-sample ARG. The recombination rate *ρ* was set to 0.166 and 11.316 in the left and right plots, respectively, such that *P* (*T*_left_ = *T*_right_) equals 0.9 or 0.1. The red diagonal identity line is plotted as a reference.

## 5. The structured coalescent

We now consider the structured coalescent, and use the notation and set-up described in Section 5.2 in Wakeley (2009). The number of demes (or sub-populations) is *D* ≥ 2, and we assume that the rate of migration for a lineage is the same between any two demes and is given by *M/*(2(*D* − 1)). We also assume that the coalescent rate for two lineages within any deme is one. We focus on moments and distributions of coalescent times for samples of size 2. Since this model is completely symmetric we only need three states: a ‘within’ state where the two lineages are in the same deme, a ‘between’ state where the lineages are in two different demes, and a ‘common ancestry’ state where the two lineages have coalesced. The last state is an absorbing state.

The ancestral process transitions from the ‘within’ state to the ‘between’ state with rate *M* because we have two lineages and each lineage can migrate to (*D* − 1) demes. The rate of coalescent is one in the ‘within’ state. The transition rate from the ‘between’ state to the ‘within’ state is *M/*(*D* − 1) because one of the two lineages has to move to exactly that deme where the other lineage is located. It is impossible to coalesce in the ‘between’ state because the lineages are in different demes. The subintensity matrix is thus given by

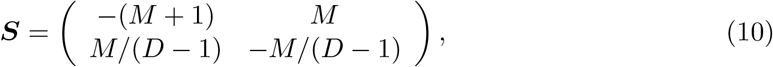

where the order of states is first ‘within’ and second ‘between’.

It is straight-forward to use **PhaseTypeR** to determine the mean and variance for a given number of demes and varying migration rate.

~~~
*R> initial_within <- c(1, 0)
R> initial_between <- c(0, 1)
R> structured_subintensity_matrix <- function(deme_number, migration_rate){
+     subintensity_matrix <- matrix(
+        c(-migration_rate-1,  migration_rate*,
*+           migration_rate/(deme_number-1), -migration_rate/(deme_number-1))*,
*+     nrow=2, ncol=2, byrow=TRUE)
+  subintensity_matrix
+ }
R> n <- 200
R> mig_rate_vec <- seq(0*.*01, 10, len=n)
R> mean_within <- rep(0, n)
R> mean_between <- rep(0, n)
R> var_within <- rep(0, n)
R> var_between <- rep(0, n)
R> for (i in 1:n){
+  structured_subint_mat <-
+     structured_subintensity_matrix(deme_number=10, mig_rate_vec[i])
+  withinPH <- PH(structured_subint_mat, initial_within)
+  mean_within[i] <- mean(withinPH)
+  var_within[i] <- var(withinPH)
+  betweenPH <- PH(structured_subint_mat, initial_between)
+  mean_between[i] <- mean(betweenPH)
+  var_between[i] <- var(betweenPH)
+ }*
~~~

The resulting plots are shown in Figure 5, and they reproduce Figure 5.1 in Wakeley (2009). We note that the mean coalescent time for two samples from the same deme is independent of the migration rate: the mean time ***e***_1_(−***S***)^−1^***e*** = *D* (recall Table 1) is the number of demes *D*. We also note that the mean and variance are substantially different for the two starting states when the migration is low, but converge when the migration rate is high. Similarly we can find the density functions for the coalescent time.

**Figure 5:**
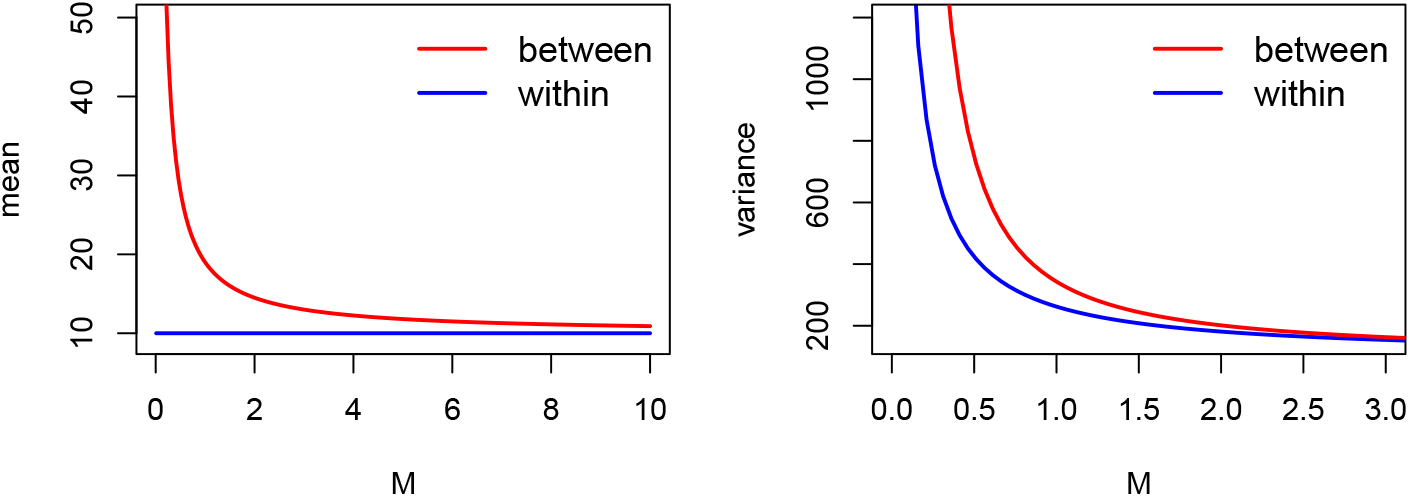
Mean (left) and variance (right) of the coalescent time across varying migration rates, with ten demes (*D* = 10) and two samples. The initial state was set to either within the same deme (blue, state 1) or between demes (red, state 2).

~~~
*R> x <- seq(0, 14, length*.*out = 100)
R> structured_subint_mat_1 <-
+    structured_subintensity_matrix(deme_number=2, migration_rate=1*.*0)
R> structured_subint_mat_2 <-
+    structured_subintensity_matrix(deme_number=10, migration_rate=1*.*0)
R> ## Initial state within:
R> withinPH_1 <- PH(structured_subint_mat_1, initial_within)
R> withinPDF_1 <- dPH(x, withinPH_1)
R> withinPH_2 <- PH(structured_subint_mat_2, initial_within)
R> withinPDF_2 <- dPH(x, withinPH_2)
R> ## Initial state between:
R> betweenPH_1 <- PH(structured_subint_mat_1, initial_between)
R> betweenPDF_1 <- dPH(x, betweenPH_1)
R> betweenPH_2 <- PH(structured_subint_mat_2, initial_between)
R> betweenPDF_2 <- dPH(x, betweenPH_2)*
~~~

In Figure 6 we show the densities of the coalescent times with fixed migration rate *M* = 1, deme number *D* = 2 or *D* = 10, and initial state either within (left plot) or between (right plot). Figure 6 in this article reproduces figures 5.2 and 5.3 in Wakeley (2009). Perhaps the most striking difference between the left and right plot is that the coalescence density is monotonocially decreasing when the initial sampling is within one deme, whereas the coalescence density is unimodal when the initial sampling is between two demes.

**Figure 6:**
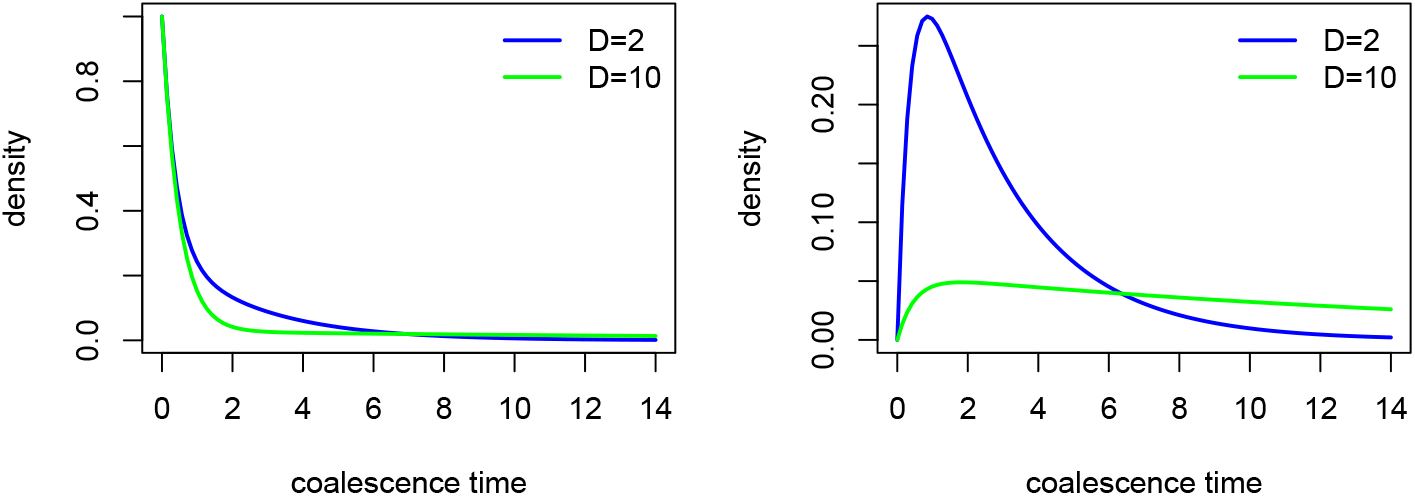
Density of the coalescent time between two samples, where the migration rate is fixed to *M* = 1 and the initial state is set to either within the same deme (left) or between demes (right). Blue and green indicate a number of demes of *D* = 2 or *D* = 10, respectively.

## 6. Conclusion, discussion and perspectives

In **PhaseTypeR** we have implemented the key characteristics and desired functions for the DPH, MDPH, PH and MPH distributions, and in this paper we have illustrated the usage in simple examples (Section 2 and Section 3) and more involved applications (Section 4 and Section 5) from population genetics. The ability to reward-transform is particularly important in population genetics, and a unique feature of **PhaseTypeR**.

We have demonstrated that phase-type theory in general and **PhaseTypeR** in particular contains the basic foundation and implementation for obtaining insight and understanding a wide range of population genetic models. In Section 2 we concentrated on the standard Kingman’s coalescent, in Section 3 on the coalescent with mutation, in Section 4 on the ancestral recombination graph, and in Section 5 on the structured coalescent. All of these models are homogeneous in time and determined by the instantaneous rate matrix and initial distribution. Other time-homogeneous population genetic models include the multiple merger coalescent (Tellier and Lemaire 2014; Freund 2021; Birkner and Blath 2021) and dormancy (Blath and Kurt 2021).

A major challenge with the coalescent models is the rapid increase in the size of the state space with the number of samples. Indeed, in the simple standard Kingman’s coalescent, the size of the state space equals the partition number from number theory (Hobolth *et al*. 2021), which increases exponentially fast in the the square root of the sample size. The instantaneous rate matrices for the coalescent models are often sparse, and with a clear structure (i.e. the number of ancestral lineages always decreases except when recombination is present). The current version of **PhaseTypeR** is not taking advantage of such special structure, but it could be important for future versions because the size of population genetic data sets are often very large.

Another extension of **PhaseTypeR** could be to allow for in-homogeneity in time. For example, Arredondo, Mourato, Nguyen, Boitard, Rodríguez, Noûs, Mazet, and Chikhi (2021) consider a structured coalescent where the number of demes is constant in time, but the migration rate has different values in epochs of time in the past. Such a model requires to paste together the probabilities from the different epochs, and intermediate epochs require the calculation of the matrix exponential (see e.g. supplementary material in Zeng, Charlesworth, and Hobolth (2021)). Recent progress for calculating the matrix exponential for large rate matrices is available in Sherlock (2021).

Applications of phase-type distributions for statistical inference is still in its infancy, but we hope that this package will fuel the development. We have demonstrated how to determine the mean and co-variance matrix for the site frequency spectrum from a coalescent model given the sample size and a set of parameters. A natural procedure for estimating the parameters of a coalescent model using phase-type theory is to match the observed and expected site frequency spectrum. Birkner and Blath (2021) describe inference methods for coalescent models with highly skewed offspring distributions using the site frequency spectrum and the methods of moments for parameter estimation.

## Supporting information

Accompanying code

## Acknowledgements

We are grateful to Emil Dare, Moisès Coll Maciá, Kasper Munch and Mikkel Heide Schierup for valuable comments and suggestions that helped improve the manuscript.

## Notes

### Competing Interest Statement

The authors have declared no competing interest.

