## Supplementary material for "PhaseTypeR: phase-type distributions in R with reward transformations and a view towards population genetics": Accompanying code

```
## Load package
# install.packages('PhaseTypeR')
library(PhaseTypeR)

##################################################
# Univariate continuous phase-type distributions #
##################################################

## Defining the distribution
subintensity_matrix <- matrix(c(-6,  6,  0,
                                0, -3,  3,
                                0,  0, -1),
                              ncol = 3, byrow = T)
initial_probabilities <- c(1, 0, 0)
T_MRCA <- PH(subintensity_matrix, initial_probabilities)
T_MRCA
```

```
## $subint_mat
##      [,1] [,2] [,3]
## [1,]   -6    6    0
## [2,]    0   -3    3
## [3,]    0    0   -1
## 
## $init_probs
##      [,1] [,2] [,3]
## [1,]    1    0    0
## 
## $defect
## [1] 0
## 
## attr(,"class")
## [1] "cont_phase_type"
```

```
## Mean and variance
mean(T_MRCA)
```

```
## [1] 1.5
```

```
var(T_MRCA)
```

```
## [1] 1.138889
```

```
## Functions for PH distributions
dPH(c(0.1, 0.5, 0.8), T_MRCA)
```

```
## [1] 0.06482665 0.48210919 0.54651397
```

```
pPH(c(0.1, 0.5, 0.8), T_MRCA)
```

```
## [1] 0.002348541 0.121417559 0.280279868
```

```
qPH(c(0.05, 0.5, 0.95), T_MRCA)
```

```
## [1] 0.3302855 1.2328314 3.5830871
```

```
set.seed(3)
rPH(3, T_MRCA)
```

```
## [1] 0.6459884 0.1019513 1.0577725
```

```
set.seed(3)
rFullPH(T_MRCA)
```

```
##   state      time
## 1     1 0.1025055
## 2     2 0.3346948
## 3     3 0.2087881
```

```
## Reward transformation
reward <- c(4, 3, 2)
T_total <- reward_phase_type(T_MRCA, reward)
T_total
```

```
## $subint_mat
##      [,1] [,2] [,3]
## [1,] -1.5  1.5  0.0
## [2,]  0.0 -1.0  1.0
## [3,]  0.0  0.0 -0.5
## 
## $init_probs
##      [,1] [,2] [,3]
## [1,]    1    0    0
## 
## $defect
## [1] 0
## 
## attr(,"class")
## [1] "cont_phase_type"
```

```
c(mean(T_total), var(T_total))
```

```
## [1] 3.666667 5.444444
```

```
# Matches general formulas from Wakeley Section 3.3:
n <- 4
c( 2*sum(1/(1:(n-1))), 4*sum(1/(1:(n-1))^2) )
```

```
## [1] 3.666667 5.444444
```

```
####################################################
# Multivariate continuous phase-type distributions #
####################################################

## Defining the distribution
subintensity_matrix <- matrix(c(-6, 6,  0,  0,
                                0, -3,  1,  2,
                                0,  0, -1,  0,
                                0,  0,  0, -1),
                              ncol = 4, byrow = T)
initial_probabilities <- c(1, 0, 0, 0)
reward_matrix <- matrix(
  c(4,2,0,1,
    0,1,2,0,
    0,0,0,1),
  nrow = 4)
L <- MPH(subintensity_matrix, initial_probabilities, reward_matrix)
L
```

```
## $subint_mat
##      [,1] [,2] [,3] [,4]
## [1,]   -6    6    0    0
## [2,]    0   -3    1    2
## [3,]    0    0   -1    0
## [4,]    0    0    0   -1
## 
## $init_probs
##      [,1] [,2] [,3] [,4]
## [1,]    1    0    0    0
## 
## $reward_mat
##      [,1] [,2] [,3]
## [1,]    4    0    0
## [2,]    2    1    0
## [3,]    0    2    0
## [4,]    1    0    1
## 
## $defect
## [1] 0
## 
## attr(,"class")
## [1] "mult_cont_phase_type"
```

```
# Calculating the variance-covariance matrix
var(L)
```

```
##            [,1]       [,2]       [,3]
## [1,]  1.7777778 -0.2222222  0.8888889
## [2,] -0.2222222  2.3333333 -0.4444444
## [3,]  0.8888889 -0.4444444  0.8888889
```

```
################################################
# Univariate discrete phase-type distributions #
################################################

## Defining the distribution
tht <- 3
p_1 <- tht/(2+tht)
p_12 <- 2/(2+tht)*tht/(1+tht)
p_2 <- tht/(1+tht)
T_mat <- matrix(c(p_1, p_12,
                  0,   p_2),
                ncol = 2, byrow = T)
init_probs <- c(p_1, p_12)
S_total <- DPH(T_mat, init_probs)
S_total
```

```
## $subint_mat
##      [,1] [,2]
## [1,]  0.6 0.30
## [2,]  0.0 0.75
## 
## $init_probs
##      [,1] [,2]
## [1,]  0.6  0.3
## 
## $defect
## [1] 0.1
## 
## attr(,"class")
## [1] "disc_phase_type"
```

```
## Mean and variance
mean(S_total)
```

```
## [1] 4.5
```

```
var(S_total)
```

```
## [1] 15.75
```

```
## Functions for DPH distributions
dDPH(c(0, 1, 2, 10), S_total)
```

```
## [1] 0.10000000 0.13500000 0.13725000 0.02573811
```

```
pDPH(c(0, 1, 2, 10), S_total)
```

```
## [1] 0.1000000 0.2350000 0.3722500 0.9191577
```

```
qDPH(c(0.05, 0.5, 0.95), S_total)
```

```
## [1]  0  4 12
```

```
set.seed(3)
rDPH(5, S_total)
```

```
## [1] 14  3  8  8 12
```

```
set.seed(3)
rFullDPH(S_total)
```

```
##   state time
## 1     1    1
## 2     2   13
```

```
## Reward transformation
T_mat <- matrix(c(p_1, p_12/2, p_12/2,
                  0,   p_2/2,  p_2/2,
                  0,   p_2/2,  p_2/2),
                ncol = 3, byrow = T)
init_probs <- c(p_1, p_12/2, p_12/2)
S_total <- DPH(T_mat, init_probs)
# Singletons
singletons <- reward_phase_type(S_total, c(1, 1, 0))
singletons
```

```
## $subint_mat
##      [,1] [,2]
## [1,]  0.6 0.24
## [2,]  0.0 0.60
## 
## $init_probs
##      [,1] [,2]
## [1,]  0.6 0.24
## 
## $defect
## [1] 0.16
## 
## attr(,"class")
## [1] "disc_phase_type"
```

```
# Doubletons
doubletons <- reward_phase_type(S_total, c(0, 0, 1))
doubletons
```

```
## $subint_mat
##      [,1]
## [1,]  0.6
## 
## $init_probs
##      [,1]
## [1,]  0.6
## 
## $defect
## [1] 0.4
## 
## attr(,"class")
## [1] "disc_phase_type"
```

```
# Mean
c(mean(singletons), mean(doubletons))
```

```
## [1] 3.0 1.5
```

```
##################################################
# Multivariate discrete phase-type distributions #
##################################################

## Defining the distribution
SFS <- MDPH(T_mat, init_probs, matrix(c(1, 1, 0, 0, 0, 1), nrow = 3))
SFS
```

```
## $subint_mat
##      [,1]  [,2]  [,3]
## [1,]  0.6 0.150 0.150
## [2,]  0.0 0.375 0.375
## [3,]  0.0 0.375 0.375
## 
## $init_probs
##      [,1] [,2] [,3]
## [1,]  0.6 0.15 0.15
## 
## $reward_mat
##      [,1] [,2]
## [1,]    1    0
## [2,]    1    0
## [3,]    0    1
## 
## $defect
## [1] 0.1
## 
## attr(,"class")
## [1] "mult_disc_phase_type"
```

```
## Variance-covariance matrix
var(SFS)
```

```
##      [,1] [,2]
## [1,] 7.50 2.25
## [2,] 2.25 3.75
```

```
#####################################
# The coalescent with recombination #
#####################################

recomb_rate <- 0.3
ARG_subint_mat <- function(recomb_rate) {
  matrix(
    c(-(1+2*recomb_rate/2), 2*recomb_rate/2,  0,             0,  0,
        1,                -(3+recomb_rate/2), recomb_rate/2, 1,  1,
        0,                  4,               -6,             1,  1,
        0,                  0,                0,            -1,  0,
        0,                  0,                0,             0, -1),
    nrow=5, byrow=TRUE)
}
subintensity_matrix <- ARG_subint_mat(recomb_rate)
initial_probabilities <- c(1, 0, 0, 0, 0)
# T_left: T_MRCA in left locus
reward_left <- c(1, 1, 1, 0, 1)
# T_right: T_MRCA in right locus
reward_right <- c(1, 1, 1, 1, 0)
# Joint distribution T_joint of T_left and T_right
T_joint <- MPH(subintensity_matrix,
               initial_probabilities,
               matrix(c(reward_left, reward_right), nrow = 5))
# Verify that phase-type result and classical formula are identical
c(var(T_joint)[1, 2],
  (recomb_rate + 18) / (recomb_rate ^ 2 + 13 * recomb_rate + 18))
```

```
## [1] 0.8321965 0.8321965
```

```
## Simulation from the joint distribution
subintensity_matrix_09 <- ARG_subint_mat(0.166)
Tab_09 <- MPH(subintensity_matrix_09, initial_probabilities,
              matrix(c(reward_left, reward_right), nrow=5))
subintensity_matrix_01 <- ARG_subint_mat(11.316)
Tab_01 <- MPH(subintensity_matrix_01, initial_probabilities,
              matrix(c(reward_left, reward_right), nrow=5))

set.seed(3)
rTab_09 <- rMPH(1000, Tab_09)
rTab_01 <- rMPH(1000, Tab_01)
cat("Empirical correlation:",cor(rTab_09[,1],rTab_09[,2]),"\n")
```

```
## Empirical correlation: 0.8987308
```

```
cat("Empirical correlation:",cor(rTab_01[,1],rTab_01[,2]),"\n")
```

```
## Empirical correlation: 0.1046957
```

```
# pdf(file="fig_simonsen_cor.pdf", width = 8, height = 4)
par(mfrow=c(1,2), par(mar=c(4.5, 4.5, 1, 1)))
{
  plot(rTab_09[,1],rTab_09[,2],pch=19,cex=0.1,
       xlim = c(0, 7), ylim = c(0, 7),
       ylab=expression(T[left]), xlab=expression(T[right]),
       panel.first = abline(a=0,b=1,col='red'))
}
{
  plot(rTab_01[,1],rTab_01[,2],pch=19,cex=0.1,
       xlim = c(0, 7), ylim = c(0, 7),
       ylab=expression(T[left]), xlab=expression(T[right]),
       panel.first = abline(a=0,b=1,col='red'))
}
```

```
# dev.off()
```

```
#############################
# The structured coalescent #
#############################

## Reproduction of Figure 5.1 in Wakeley (2009)
initial_within <- c(1, 0)
initial_between <- c(0, 1)
structured_subintensity_matrix <- function(deme_number, migration_rate){
  subintensity_matrix <- matrix(
    c(-migration_rate-1,                migration_rate,
      migration_rate/(deme_number-1), -migration_rate/(deme_number-1)),
    nrow=2, ncol=2, byrow=TRUE)
  subintensity_matrix
}
n <- 200
mig_rate_vec <- seq(0.01, 10, len=n)
mean_within <- rep(0, n)
mean_between <- rep(0, n)
var_within <- rep(0, n)
var_between <- rep(0, n)
for (i in 1:n){
  structured_subint_mat <-
    structured_subintensity_matrix(deme_number=10, mig_rate_vec[i])
  withinPH <- PH(structured_subint_mat, initial_within)
  mean_within[i] <- mean(withinPH)
  var_within[i] <- var(withinPH)
  betweenPH <- PH(structured_subint_mat, initial_between)
  mean_between[i] <- mean(betweenPH)
  var_between[i] <- var(betweenPH)
}
```

```
# pdf(file="fig_5_1.pdf", width = 8, height = 3)
par(mfrow=c(1,2), par(mar=c(4.5, 4.5, 1, 1)))
## Plot mean values of coalescent time between pairs of lineages
## when the beginning state is either within or between,
## the number of demes is D=10, and for various migration rates.
{
  plot(mig_rate_vec, mean_within, type="l",
       # main="Mean time to coalescence for D=10 and varying M",
       main=NULL,
       col="blue", lwd=2, ylim=c(9,50), ylab="mean", xlab="M")
  points(mig_rate_vec,mean_between, type="l", col="red", lwd=2)
  legend("topright", c("between", "within"), lty=1, lwd=2,
         col=c("red", "blue"), bty="n", cex=1.2)
}
## Variance of coalescent times between pairs of lineages
{
  plot(mig_rate_vec, var_within, type="l", xlim=c(0, 3),
       # main="Variance of coalescent time for D=10 and varying M",
       main=NULL,
       col="blue", lwd=2, ylim=c(150,1200), ylab="variance", xlab="M")
  points(mig_rate_vec,var_between,type="l", col="red", lwd=2)
  legend("topright", c("between", "within"), lty=1, lwd=2,
         col=c("red", "blue"), bty="n", cex=1.2)
}
```

```
# dev.off()
```

```
## Reproduction of Figure 5.3 in Wakeley (2009)
## (note: Figure 5.2 is a subset of Figure 5.3)

# Left panel in Figure 5.3:
x <- seq(0,14, length.out = 100)
structured_subint_mat_1 <-
  structured_subintensity_matrix(deme_number=2, migration_rate=1.0)
withinPH_1 <- PH(structured_subint_mat_1, initial_within)
withinPDF_1 <- dPH(x, withinPH_1)
structured_subint_mat_2 <-
  structured_subintensity_matrix(deme_number=10, migration_rate=1.0)
withinPH_2 <- PH(structured_subint_mat_2, initial_within)
withinPDF_2 <- dPH(x, withinPH_2)

# Right panel in Figure 5.3:
betweenPH_1 <- PH(structured_subint_mat_1, initial_between)
betweenPDF_1 <- dPH(x, betweenPH_1)
betweenPH_2 <- PH(structured_subint_mat_2, initial_between)
betweenPDF_2 <- dPH(x, betweenPH_2)
```

```
# pdf(file="fig_5_3.pdf", width = 8, height = 3)
par(mfrow=c(1,2), par(mar=c(4.5, 4.5, 1, 1)))
{
  plot(x, withinPDF_1, type="l",
       col="blue", lwd=2, ylab="density", xlab="coalescence time",
       # main="Within probability densities when migration rate is 1.0"
       main=NULL
  )
  points(x, withinPDF_2, type="l", col="green", lwd=2)
  legend("topright", c("D=2", "D=10"), lty=1, lwd=2,
         col=c("blue", "green"), bty="n")
}

{
  plot(x, betweenPDF_1, type="l",
       col="blue", lwd=2, ylab="density", xlab="coalescence time",
       # main="Between probability densities when migration rate is 1.0"
       main=NULL
  )
  points(x, betweenPDF_2, type="l", col="green", lwd=2)
  legend("topright", c("D=2", "D=10"), lty=1, lwd=2,
         col=c("blue", "green"), bty="n")
}
```

```
# dev.off()
```
